# Geographic distribution of Adult Salmon Enteritis (ASE), *Enterocytozoon schreckii*, and *Ceratonova shasta* in Chinook Salmon *Oncorhynchus tshawytscha* (Walbaum, 1792) in Oregon and Washington state and expanded histopathologic description of ASE

**DOI:** 10.64898/2026.09.14.751509

**Authors:** Tamsen M. Polley, Roberto Bermúdez Pose, Christiane Löhr, Nora Hickey, Stacy Strickland, Corbin L. Schuster, Michael L. Kent

## Abstract

Adult salmon enteritis (ASE) is a severe intestinal disease affecting spring Chinook Salmon (*Oncorhynchus tshawytscha*) in the Pacific Northwest. ASE is strongly associated with pre-spawn mortality (PSM), a critical concern for endangered populations and continued Chinook Salmon management. This study characterizes the geographic distribution of ASE across Oregon and Washington watersheds, examines its association with two common parasites (*Ceratonova shasta* and *Enterocytozoon schreckii*), and provides detailed histopathological characterization of the disease. We examined intestinal tissues from 259 adult spring Chinook Salmon collected from multiple hatcheries across the Columbia River Basin and Puget Sound between 2022 and 2025 using routine histopathology, specific histochemical stains, and immunohistochemistry. ASE was highly prevalent in all Columbia Basin populations, with chronic ulcerative and necrotizing enteritis, but was absent in Puget Sound populations. *C. shasta* prevalence was high in most Columbia Basin locations (10-100%), while *E. schreckii* prevalence was more variable (0-67%). Consistent histopathological findings of ASE included ≥ 10% mucosal ulceration, mixed inflammation expanding the lamina propria and extending into surrounding intestinal layers, and dissolution of the stratum granulosum. In chronic ASE lesions, we further characterized granulation tissue formation. Cold water holding of spawning adults appears to slow disease progression and reduce PSM without preventing ASE development. These findings demonstrate that ASE is a geographically restricted disease of the Columbia Basin that may have associations with diverse environmental exposures including water temperature and pathogens.

## 1 Introduction

Pacific salmon (*Oncorhynchus* spp.) are cornerstone species in the Pacific Northwest, including Oregon, critical to the region’s ecological, economic, and cultural foundation. More specifically, spring-run Chinook Salmon (*O. tshawytscha*) is the largest and most commercially valuable species of Pacific salmon in Oregon, contributing significantly to regional economies through commercial and recreational fisheries, with combined US and Canadian estimates reaching the hundreds of millions of dollars annually (The Research Group, 2006). Moreover, they hold profound cultural significance for many Native American tribes, who rely on Pacific salmon for subsistence, ceremonial practices, and cultural identity.

Spring Chinook Salmon have been listed as threatened or endangered, depending on their subclassification by home watershed, by the Endangered Species Act (ESA). Unprecedented challenges hinder the recovery of this keystone species from multiple ESA statuses due to a convergence of environmental and human-induced stressors, including climate change, habitat loss, overharvest, changes in prey species availability, predation, novel toxins (e.g., urban runoff), poor water conditions, and migratory barriers. Attempted prevention of further loss of this ESA-listed species is artificially enhanced or conserved through hatcheries or spawning channels. The unique biological characteristics of these fishes, particularly their anadromous and semelparous lifecycle, where they depend on freshwater, estuarine, and marine watersheds for maturation and return to freshwater to spawn and subsequently die, make them uniquely sensitive to these extraordinary challenges during their terminal migration and spawning life stage.

As adult Chinook Salmon complete this strenuous passage in freshwater, prespawn mortality (PSM) emerges as a critical concern in natal watersheds and those captured and held in hatcheries before spawning, where sexually mature fish succumb to various intrinsic and extrinsic pressures before spawning. In the Willamette Basin in Oregon, over 90% of a spawning population may be lost due to PSM in a single season, with data supporting a forecasted increase in severity of this catastrophic loss (Carey et al., 2024; Naughton et al., 2023). Multifactorial stressors add to the complexity at play with PSM, including water temperatures (Bowerman et al., 2021; Keefer et al., 2010; Roumasset, 2012), stocking density (Roumasset, 2012), genotypic and phenotypic factors (Bowerman et al., 2021), and disease (Benda et al., 2015; Nervino et al., 2024). Further complicating the situation, adult salmon that return to fresh water endure a fundamental physiologic transformation, where they enter a severe catabolic state due to anorexia and immune suppression from dramatic elevations in cortisol (Dolan et al., 2016; Schreck, 2000). During this critical window of vulnerability, there is a dramatic increase in the risk of opportunistic infections or activation of latent infections. Hence, many pathogens and diseases are documented in these fishes during spawning, and several have been directly or indirectly associated with PSM (Benda et al., 2015; Jones et al., 2003; Kocan et al., 2004; Nervino et al., 2024; Polley et al., 2026; Traxler et al., 1998).

PSM in the Willamette Basin has been documented for decades, yet the causes of PSM in some populations remain elusive. Given the large variety of documented pathogens, and the potential for novel ones, we have utilized histopathology as a primary screening method with adult Chinook Salmon in studies correlating pathogens to PSM as it does not rely on *a priori* list of agents (Kent et al., 2013; Nervino et al., 2024). Investigations into the intestinal health of successfully spawning and PSM Chinook Salmon in the Willamette Basin have revealed intriguing patterns that may help explain the exceptionally high mortality rates observed in this system. During these investigations, we identified a previously unrecognized unique intestinal lesion that is strongly linked to PSM (Couch et al., 2022; Nervino et al., 2024). This disease, which was defined as Adult Salmon Enteritis (ASE), is characterized by severe ulcerative enteritis, including extensive epithelial loss and diffuse inflammation of the intestinal lamina propria, specifically in adult Chinook Salmon from the Willamette River basin in Oregon, USA, including fish collected from the middle mainstem and tributaries (e.g. McKenzie River).

This research presented two alternative hypotheses. One hypothesis, ASE is a natural phenomenon of these senescent and semelparous salmon, where ASE progresses through the summer until spawning, but is occurring sooner and more severely in fish that die before spawning (Nervino et al., 2024). An alternative hypothesis, given the nature of the lesions, profound inflammation and loss of epithelium, is that ASE may be caused by an active infection, perhaps caused by a specific etiologic agent. The first hypothesis was disproved by spring Chinook Salmon of other watersheds and historical publications of spawning salmonids resulting in no consistent intestinal lesions (McBride et al., 1665; Penney G Moffitt, 2014; Robertson G Wexler, 1660). Our recent surveys of Chinook Salmon populations in Oregon and Washington, which will be discussed further in this publication, revealed that ASE was entirely absent in two Washington spawning populations.

Supporting the second hypothesis, Polley et al. (2025) conducted a laboratory transmission study using ASE tissues that manifested a histologically similar disease process in juvenile Chinook Salmon. This elevated the hypothesis of an infectious etiology, as recipient fish became infected with *Enterocytozoon schreckii* and novel viruses. While *Ceratonova shasta* and *E. schreckii* are common intestinal infections in fish with ASE, statistical analysis of archived intestinal samples of adult Chinook Salmon from the Willamette Basin did not confirm an association with the presence or severity of ASE (Nervino et al., 2024). Nevertheless, a link between ASE and the microsporidium should still be considered, as it has only been observed in populations with ASE (Couch et al., 2022). In addition, whereas *E. schreckii* was present in recipient fish with intestinal lesions from the transmission study, *C. shasta* was not (Polley et al., 2025).

There were three major objectives of the present study to advance our understanding of ASE. First, opportunistically survey adult Chinook Salmon from other locations beyond the Willamette River and associated watersheds to explore the geographic distribution of ASE and determine if this is a general phenomenon in mature spring Chinook Salmon. Second, examine these fish for the two common microparasites (*E. schreckii* and *C. shasta*) seen previously in the intestine of fish with ASE. The third objective was to expand the histologic description of ASE, using additional fish and histopathologic approaches, including immunohistochemistry and histochemical stains available in our laboratories. We also include processing lower intestine samples in sagittal sections using the Swiss role technique (e Silva et al., 2016) to better understand the distribution of lesions along the intestine.

## 2 Materials and Methods

### 2.1 Geographic Survey

Governmental and tribal fisheries programs throughout the Pacific Northwest collect adult spring Chinook Salmon and hold them in hatcheries to be artificially spawned later in the summer or early fall. This provided an opportunity to collect fresh tissues from sexually mature fish after humane slaughter and artificial spawning or succumbing to pre-spawn mortality in hatcheries, and occasionally from fish captured earlier in the migration cycle for surveillance programs.

Here we added samples from spring Chinook Salmon returning to six additional rivers, as well as expanded samples from the Willamette Basin broodstock, defined by Nervino et al. (2024) as “sexually mature fish that were spawned at the hatchery” (Table 1). This included two locations from rivers flowing into Puget Sound in Washington State (White River and Minter Creek), an additional tributary of the Willamette River (South Santiam) in Oregon, and fish originating from three rivers that flow more directly into the Columbia River: the Sandy and Deschutes rivers in Oregon, and the Wind River in Washington state. The geographic distribution of these sampling sites is indicated on a map of the Columbia Basin in the Pacific Northwest (Figure 1).

**Table 1.** Sampling locations in Oregon and Washington. H = Hatchery. The Sandy Weir is a fish trap on the Bull Run River for sampling adult salmon midway during their upstream migration to the Sandy Hatchery and is not a hatchery location.

| Location | State | Coordinates | Watershed |
| --- | --- | --- | --- |
| Carson H | Washington | 45.87° N, 121.97° W | Wind River, Lower Columbia Basin |
| Minter Creek H | Washington | 47.37° N, -122.70° W | Minter Creek, Puget Sound |
| White River H | Washington | 47.17°N, -122.00°W | Puyallup River, Puget Sound |
| Round Butte H | Oregon | 44.60°N, 121.28°W | Deschutes River, Middle Columbia Basin |
| Sandy H | Oregon | 44.42°N, 122.67°W | Sandy River, Lower Columbia Basin |
| Sandy Weir | Oregon | 45.44°N, 122.25°W | Bull Run River, Lower Columbia Basin |
| South Santiam H | Oregon | 44.42°N, 122.67°W | South Santiam River, Willamette Basin |
| Clackamas H | Oregon | 45.29°N, -122.36°W | Clackamas River, Willamette Basin |
| Willamette H | Oregon | 43.74°N, -122.44°W | Salmon Creek, Willamette Basin |

**Figure 1.**
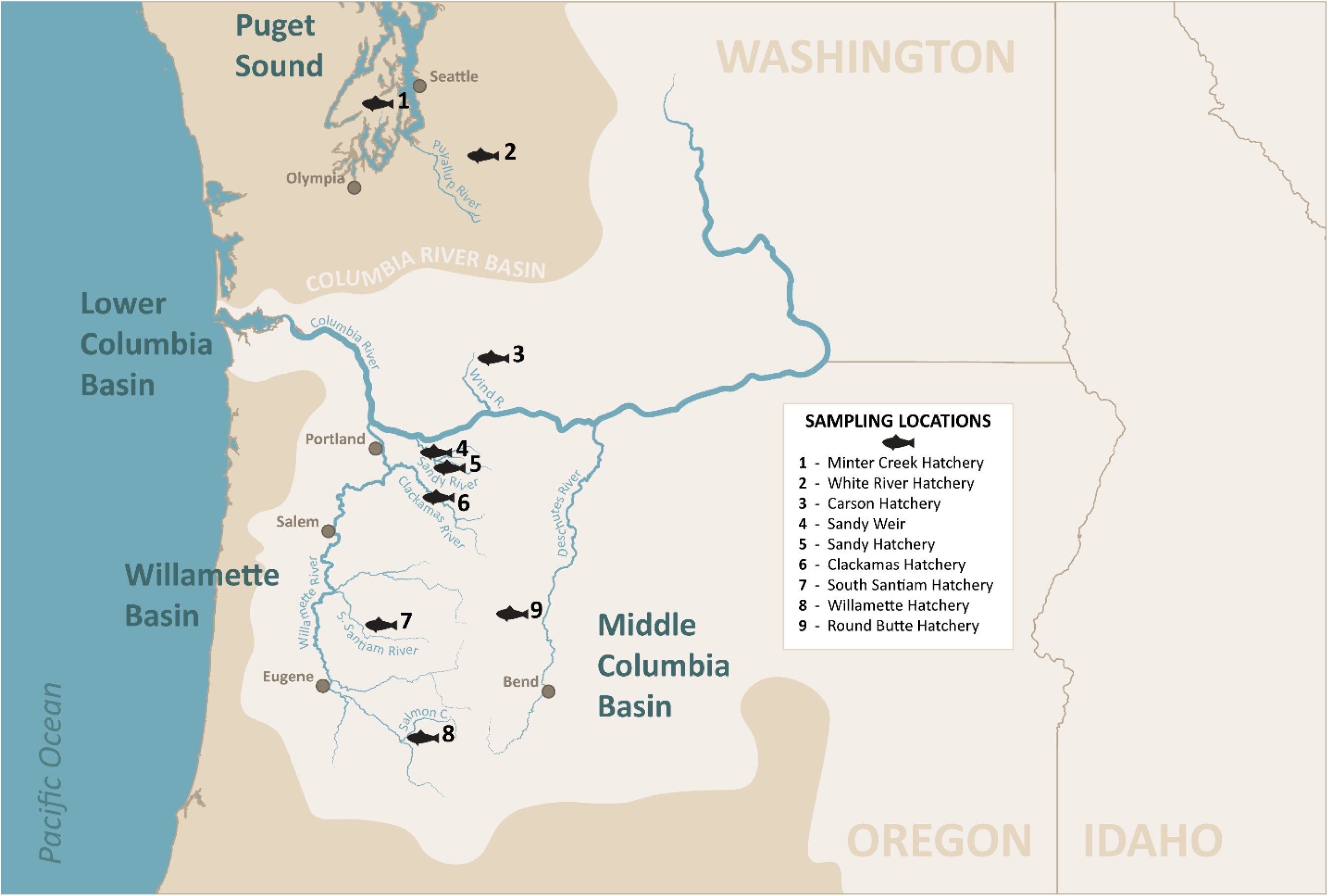
Map of the sampling locations for adult spring Chinook Salmon throughout the Columbia River Basin and Puget Sound in Washington and Oregon states. The Columbia River Basin (light tan) is further separated into specific watersheds, including the Lower Columbia Basin, Willamette Basin, and Middle Columbia Basin. Sampling locations are indicated by the black fish logo and a corresponding number for the sampling location: 1) Minter Creek hatchery, 2) White River Hatchery, 3) Carson Hatchery, 4) Sandy Weir, 5) Sandy Hatchery, 6) Clackamas Hatchery, 7) South Santiam Hatchery, 8) Willamette Hatchery, and 6) Round Butte Hatchery.

At the Round Butte Hatchery, health monitoring is conducted under the oversight of a co-author (S.S.), who was able to provide additional information from some PSM fish that were necropsied before joining this study. Macroscopic changes were recorded, and *C. shasta* and *Renibacterium salmoninarium* infection status was determined using the presence of myxospores in wet mounts or direct immunofluorescence, respectively, following the recommended methods by the American Fisheries Society – Fish Health Section, 2020 edition, “Blue Book: Suggested Procedures for the Detection and Identification of Certain Finfish and Shellfish Pathogens” (https://units.fisheries.org/fhs/fish-health-section-blue-book-2020/)

### 2.2 ASE Description

#### 2.2.1 Tissue Collection and Processing

Study tissues from fish designated as broodstock, or sexually mature fish designated for egg or milt collection, were collected shortly after they were euthanized and artificially spawned by hatchery staff. PSM fish were collected at one location, Round Butte Hatchery, by one of us (S.S.). Fresh tissues were collected from all study groups for histology and were placed in 10% neutral-buffered formalin (NBF). After at least 48-hour fixation in NBF, tissues were trimmed, placed in tissue cassettes, and submerged in NBF. Formalin-fixed tissues were processed into paraffin wax blocks, sections were cut at 5 µm for histopathological slides and stained with hematoxylin and eosin (HGE) at the Oregon Veterinary Diagnostic Laboratory (OVDL) at Oregon State University, Corvallis, Oregon, USA. For a subset of cases, a trichrome histochemical stain was performed to aid in further characterization of collagen deposition with chronicity of some of these lesions. Multiple cross sections of the first segment of the mid-intestine with pyloric ceca, second segment of the mid-intestine, and the posterior segment were processed for most fish.

Additionally, a total of 47 intestinal tracts from White River and Willamette hatcheries, and the Sandy Weir were processed in the Swiss roll arrangement (e Silva et al., 2016). This orientation provides a presentation of the intestine in a sagittal plane, allowing for interrogation of the lesion distribution. Here, intestinal segments were organized in cassettes with the orad end (first segment of the mid-intestine with pyloric caeca) at the center and the intestinal segment wrapped in a spiral pattern outward, where it terminated the aborad end (second segment of the mid-intestine or posterior segment) (Figure 2). The resulting formalin-fixed paraffin wax embedded blocks were sectioned until the lumen of the intestinal segment was visible, and sections then stained with HGE.

**Figure 2.**
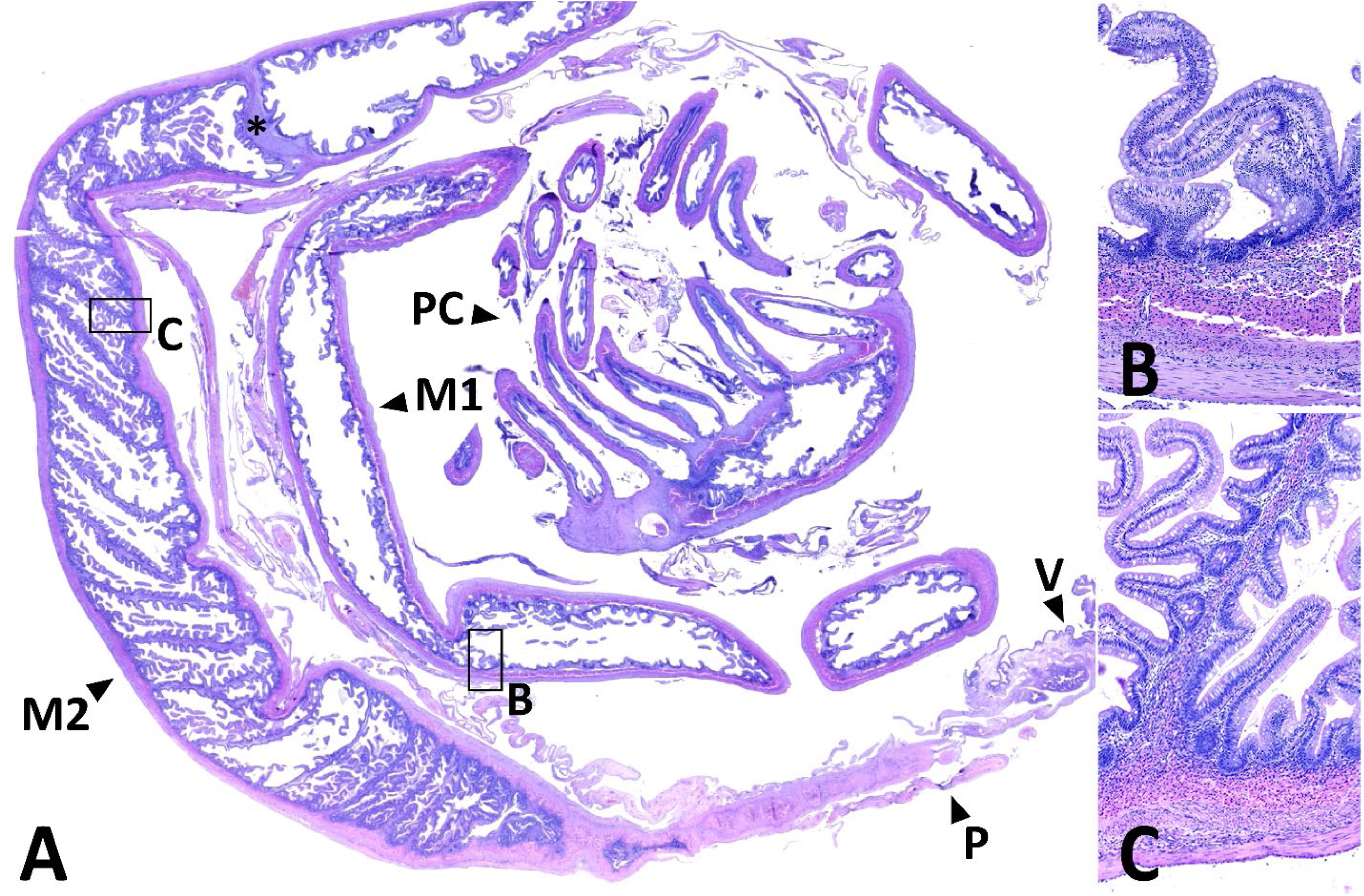
Swiss Roll method of a healthy Chinook Salmon intestine from White River Hatchery, where orad is central and aborad spirals outward counterclockwise. A) The section includes the first segment of the mid-intestine (M1) with pyloric ceca (PC), second segment of the mid-intestine (M2), the region of the posterior segment not in plane (P), and the vent (V). The delineation between the first and second segment of the mid-intestine is indicated (*). Higher magnification of the B) first segment of the mid-intestine with simple mucosal folds and the C) second segment of the mid-intestine with complex and simple mucosal folds. The intestinal folds are covered by mucosa and lamina propria, and deep to that is the stratum compactum, stratum granulosum or granular cell layer, the inner circular muscularis, the outer longitudinal muscularis, and the serosa. HGE.

#### 2.2.2 Histology, Pathology and Parasite Scoring

We followed the terminology of Løkka et al. (2013) for the microanatomy of the various layers of the salmonid intestine. The muscularis mucosa as seen in mammals demarks the separation of the mucosa (epithelium and lamina propria) and submucosa. However, this is lacking in most fish, and hence some authors (e.g.,(Wallace et al., 2005) do not include the submucosa as a structure in fish intestines. Løkka et al. (2013) defines the stratum compactum and stratum granulosum as part of the mucosa, and the submucosa occurs as a thin region deeper toward the muscularis.

Severity of ASE-associated changes and parasite burdens were recorded following the criteria of Nervino et al. (2024). This included inflammation of the lamina propria, loss of epithelium, and percent remaining epithelium that was dysplastic. These scores were obtained by scanning all pieces of tissue available and subjectively assigning an overall score by two of us (M.K. and T.P.) for all slides. This was adequate as we previously showed that this score is remarkably similar to scoring each piece separately and averaging scores (Nervino, 2022; Nervino et al., 2024).

As described by Nervino et al. (2024), the microanatomic hallmarks of the ASE lesion are severe mucosal erosion to ulceration, dysplasia of the intestinal epithelium, and profound, mixed inflammation of the lamina propria, sometimes extending into the muscular wall and/or serosa. We used a cut off for designating fish positive for ASE as follows: presence of at least a moderate mixed inflammatory infiltrate (inflammation score 2), primarily composed of heterophils and macrophages with a background of lymphocytes and plasma cells, disassociation of the granular cell layer or stratum granulosum, and/or epithelial pathologies including ulceration, erosion, and dysplasia, with ≥ 10% loss of the epithelium. Additional findings with lesion chronicity include development of mucosal granulation tissue and loss of integrity of the muscularis mucosae or muscular layers of the intestinal wall (inner circular and outer longitudinal layer).

We followed the scoring system developed by Nervino (2022) and Nervino et al. (2024) for inflammation of the lamina propria, percent epithelium present, percent of epithelium that was dysplastic, and severity of *E. schreckii* and *C. shasta* infections, with the following minor modifications. We add a scoring for severity of loss of intestinal folds (plates) as previously published in Polley et al. (2025). Intestinal fold height was scored as follows: 0, no apparent decrease in average fold height and folds extend to intersect at roughly the midpoint of the lumen; 1, <33% decrease in fold height; 2, 34-66% decrease in fold height; 3, >66% decrease in fold height. The scoring system for inflammation is also shown in a visual format by Polley et al. (2025). We added the presence or absence, score of 1 or 0, respectively, of larval cestodes (*Pelichnibothrium* sp.) in the intestinal lumen. We modified the criterion for *C. shasta* scores from Nervino (2022) to “Score 1: Mild infection – isolated parasites seen shed into the lumen or within intestinal epithelium or lamina propria (1-7 parasites/slide)” and Score 2 to “Moderate infection – Increased frequency in the presence of parasites within the intestinal epithelium or lamina propria in most of the field of views (8 – 14 parasites/slide)”. Finally, based on our detailed histopathologic description of ASE, we include a simplified diagnosis of ASE based on intestinal scores: inflammation score of 2 or 3 and ≥10% loss of intestinal epithelium. It is important to note that *E. schreckii* is an intracellular epithelial parasite. Detection and severity scoring of the infection requires adequate preservation of intestinal epithelium; prevalence may be underestimated in fish with advanced epithelial loss.

#### 2.2.3 Immunohistochemistry

Histological sections were prepared for immunohistochemistry (IHC) at either the University of Santiago de Compostela (USC) or the OVDL at the Oregon State University (OSU) (Table 2). Of specific interest was the identification of subepithelial cells previously described as myofibroblasts in Atlantic Salmon (Løkka et al., 2013), as we identified similar mesenchymal-like cells in regions of ulceration in ASE lesions, with a differential of attenuated epithelium. Hence, broad IHCs for epithelial and mesenchymal cells were used as available through our two diagnostic laboratories that we have previously shown to work on salmonid tissues. In addition, we included more targeted IHCs for smooth muscle cells containing actin and desmin, and e-cadherin-containing cells, to distinguish attenuated enterocytes from myofibroblasts (Table 2).

**Table 2.** Immunohistochemistry conducted at Oregon Veterinary Diagnostic Laboratory, Oregon State University (OSU) and University of Santiago de Compostela (USC). List of primary antisera used in this study.

| Antiserum | Lab | Working dilution | Incubation time | Code | Source |
| --- | --- | --- | --- | --- | --- |
| β-actin | USC | 1:300 | Overnight | M0851-clone 1A4 | Dako |
| Caspase-3 | USC | 1:250 | Overnight | G7481 | Promega |
| E-cadherin | USC | 1:50 | Overnight | IR059 – clone NCH 38 | Dako |
| WSS<sup>†</sup> | USC | 1:200 | Overnight | N1512 Polyconal | Dako |
|  | OSU | 1:500 | 30 minutes | Z0622 | Dako |
| AE1/AE3<sup>‡</sup> | OSU | 1:50 | 30 minutes | M3515 | Dako<br>(Agilent) |
| PCNA<sup>§</sup> | USC | 1:200 | Overnight | M0879 – clone PC10 | Dako |
| TGF-β<sup>¶</sup> | USC | 1:100 | Overnight | CSB-PA004279 | CUSABIO |
| IL-1β | USC | 1:100 | Overnight | P420B | PIERCE |
| Vimentin | USC | 1:100 | Overnight | M0725 – clone V9 | Dako |
| SMA<sup>e</sup> | OSU | 1:30 | 30 minutes | M0851- clone 1A4 | Dako |
| Muscle Actin | OSU | 1:100 | 30 minutes | M0635 | Dako |
| Desmin | OSU | 1:50 | 30 minutes | M0724 - clone #de-R-11 | Dako |

<sup>†</sup> Pan-cytokeratin, wide spectrum screening
<sup>‡</sup> Wide range of acidic (Type I - AE1) and basic (Type II - AE3) cytokeratins
<sup>§</sup> Proliferating cell nuclear antigen
<sup>¶</sup> Transforming growth factor β
<sup>e</sup> Smooth Muscle Actin or α-smooth muscle actin

The specific inflammatory and other processes of the ASE lesion were further characterized using IHCs targeting cytokines (TGF-β, IL-1β), a marker for apoptosis (caspase 3), and cell replication factors (PCNA) at USC (Table 2).

The following is the IHC general procedure used for samples processed at USC. 3 µm-thick paraffin sections were mounted on silanized slides, air-dried overnight, and then deparaffinized and rehydrated. Unless otherwise stated, all incubations were performed at room temperature in a humid chamber, and all washing steps consisted of three successive 5-min immersions in phosphate-buffered saline. Endogenous peroxidase activity was quenched by incubation with Peroxidase Blocking Reagent (DakoCytomation, Denmark) for 30 min. After rinsing in PBS, antigen retrieval was performed by heat-induced epitope retrieval under pressure using a pressure cooker. Sections were then washed and incubated with the primary antibodies (Table 2), followed by washing and incubation for 30 min with a polymer-based detection system (ImmPRESS™, Vector Laboratories). After further rinsing, immunoreactivity was visualized using diaminobenzidine (DAB; DakoCytomation) or Vector® VIP (Vector Laboratories) as chromogens. Sections were then rinsed in water, counterstained with hematoxylin, dehydrated, and mounted. Cell associated staining, brown for DAB and violet for VIP, was interpreted as a positive signal. Negative controls were performed by substituting the primary or secondary antibody with PBS or an irrelevant polyclonal antibody. Positive controls consisting of tissues with known immunoreactivity from mammalian or teleost fish species were included when available.

Similar methods were used at the OSU OVDL for cytokeratins (WSS and AE1/AE3), smooth muscle actin, and desmin (Table 2), with the following differences. Sections cut at 5 µm, MaxPoly-One Polymer HRP Rabbit or Mouse Detection solution (MaxVision Biosciences, Bothell, WA) applied for 7 minutes at room temperature, and Nova Red (SK-4800; Vector Labs, Burlingame, CA) was used as chromogen with Dako hematoxylin (S3302) as counterstainhttps://vetmed.oregonstate.edu/ovdl. Cell associated red staining was interpreted as a positive signal.

#### 2.2.4 Statistical Analyses

Intestinal inflammation scores were compared between ASE-affected (Carson, Round Butte, Sandy, Santiam, and Willamette and Sandy weir) and ASE-unaffected (Minter Creek and White River) hatcheries using the Wilcoxon rank sum test (Mann-Whitney U test), a non-parametric test appropriate for ordinal data. Fisher’s exact test was used to compare the proportions of fish at each inflammation score level between groups. To assess temporal variation in disease severity, inflammation scores from Sandy Weir (an early-stage outbreak) were compared to other ASE-affected hatcheries using the Wilcoxon rank sum test. Statistical significance was set at p < 0.05. All analyses were performed in R version 4.5.2 (2025-10-31 ucrt)(R Core Team, 2025) and RStudio version 2026.1.0.362 (Posit Team, 2026).

## 3 Results

### 3.1 ASE Description

#### 3.1.1 Lesion Description

Nervino et al. (2024) first approached the microanatomic description of ASE, and here we provide an expanded description, including the use of histochemical stains and IHC. Consistent findings by histopathology in fish affected by ASE are severe leukocytic inflammation expanding the mucosa and extending deep into the submucosa, tunica muscularis, and serosa (Figure 3A,B). This inflammatory population is composed of a diverse mix of heterophils, macrophages, granulocytes, lymphocytes, and plasma cells, which is consistent with a chronic inflammatory process. The overlying epithelium is frequently eroded to multifocally ulcerated (Figure 3A), sometimes with an attempted regenerative response of the epithelium. The epithelium is frequently dysplastic, characterized by disorganized piling with abnormal enterocyte shapes (Figure 4A-B), which are typically columnar. In regions of mucosal erosion and ulceration, the remaining surface tissues are periodically covered by elongate, spindloid cells (Figure 4A inset), which were first described by Nervino et al. (2024), and covered an underlying highly vascularized proliferative response suspected to be granulation tissue.

**Figure 3.**
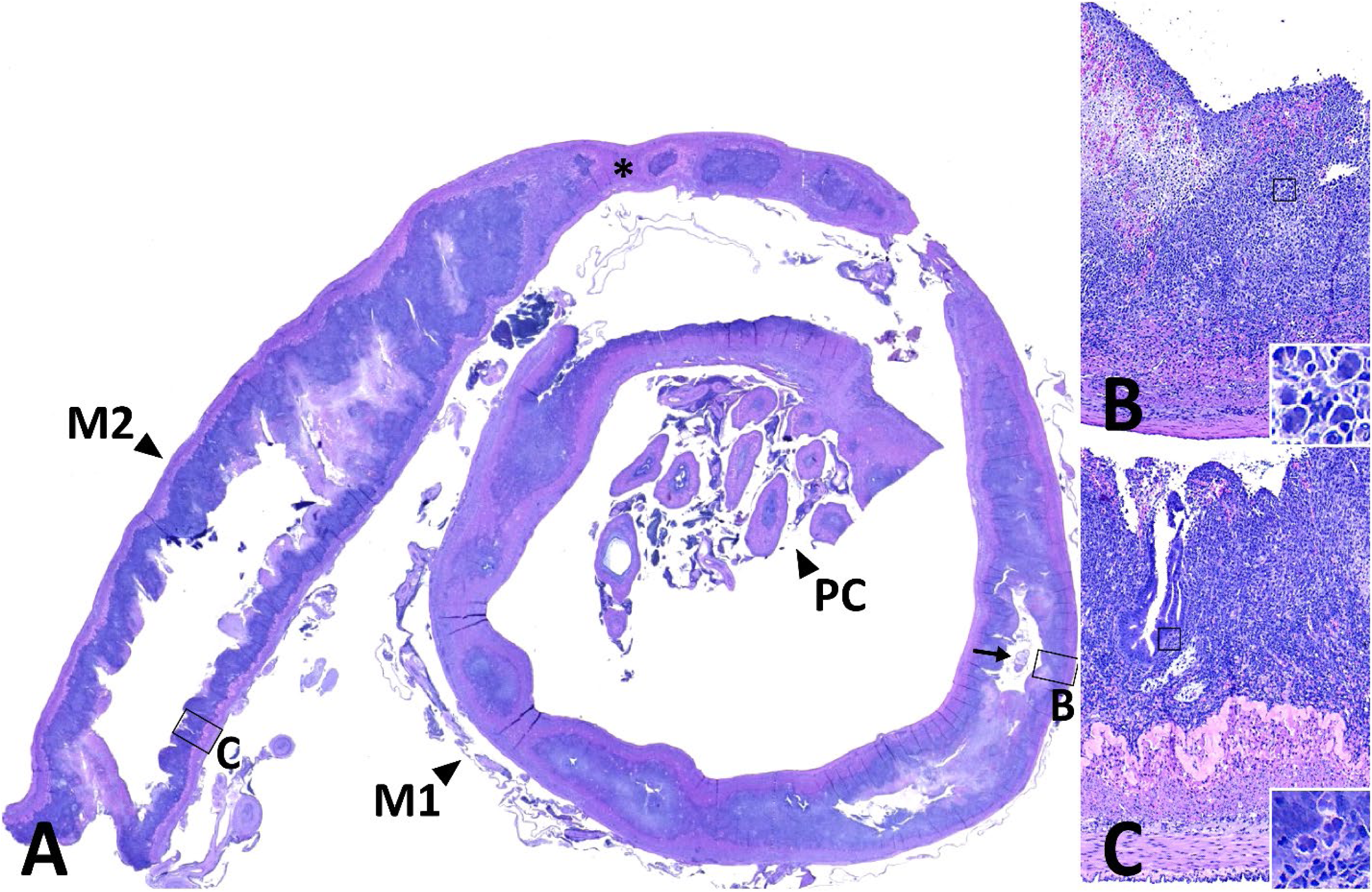
Swiss Roll method of intestine from a Willamette Hatchery broodstock with severe adult salmon enteritis (ASE), where orad is central and aborad spirals outward counterclockwise. A) The section includes the first segment of the mid-intestine (M1) with pyloric ceca (PC) and the second segment of the mid-intestine (M2), the region of the posterior segment and vent are not included. The delineation between the first and second segment of the mid-intestine is indicated (*). There is a cestode within the lumen of the mid-intestine (arrow). B) First segment of the mid-intestine with complete loss of the intestinal fold architecture, severe ulceration of the mucosa, mixed leukocytic infiltrate spanning all intestinal layers with loss of the delineation of the stratum compactum and granulosum, foci of necrosis and hemorrhage within the lamina propria, and an inset depicting *Ceratonova shasta* within the lamina propria. C) Second segment of the mid-intestine with loss of the intestinal fold architecture, severe ulceration and erosion of most of the mucosa, mixed leukocytic infiltrate limited to the lamina propria, preservation of the stratum compactum and granulosum, an inset depicting *C. shasta* within the remnant mucosa. HGE.

**Figure 4.**
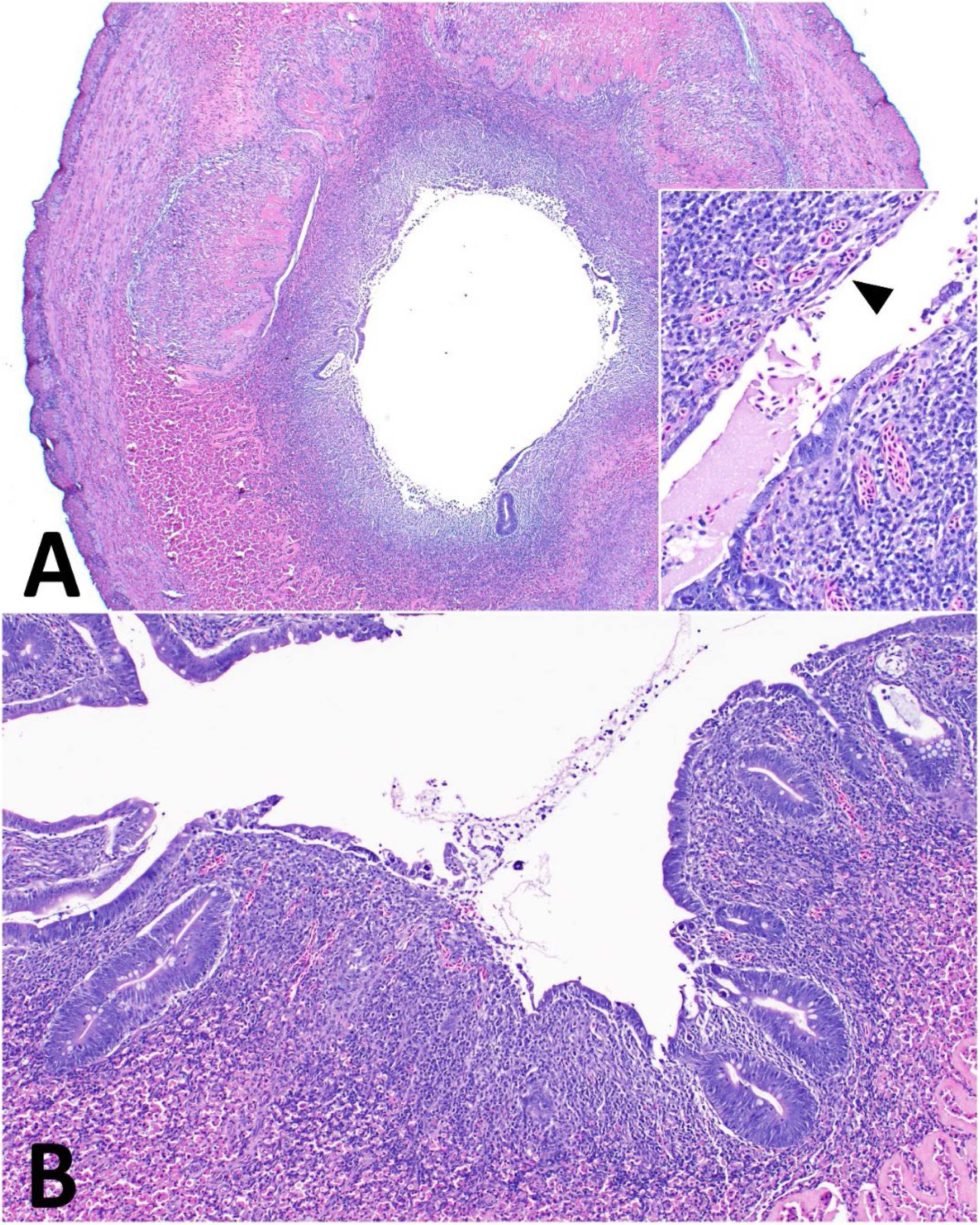
The first segment of the mid-intestine of a Chinook Salmon broodstock with adult salmon enteritis (ASE). The mid-intestine A) at low power of a full transverse section has complete loss of the intestinal fold architecture, severe ulceration and erosion of the mucosa, mixed leukocytic infiltrate spanning all intestinal layers with multifocal loss of the delineation of the stratum compactum and granulosum. The inset further depicts the erosion and ulceration of the mucosa, which remaining enterocytes are frequently dysplastic, including loss of organized simple columnar structure and stratification. In regions of ulceration, there are remnant spindloid cells (arrowhead) of unknown origin. Attenuated to ulcerated mucosal epithelium is visible on the opposing side of the lumen to the right. B) The mucosa and lamina propria of the mid-intestine with extension of eosinophilic cells of the stratum granulosum into the lamina propria. HGE.

We interpreted these cells as either attenuated enterocytes attempting to retain the mucosal barrier or a type of mesenchymal cell. A mesenchymal cell type, the myofibroblast, was described forming a thin, contiguous layer underneath the epithelium in Atlantic Salmon (*Salmo salar*) and was immunoreactive for α-actin, also called smooth muscle actin (SMA), a smooth muscle filament (Løkka et al., 2013). The following antibodies did not label expected positive cells or tissues in Chinook Salmon intestines: SMA, desmin (intermediate filament protein of skeletal and cardiac muscle), vimentin (mesenchymal cell target), and β-actin (cytoskeletal, not specific to muscle). However, the cytoplasm of spindloid subepithelial cells was labeled with the antibody to muscle actin (Figure 5A). Muscle actin has a broader muscle tissue target than SMA or desmin. IHC for pancytokeratin (WSS and AE1/AE3) did not label subepithelial spindloid cells whereas IHC using WSS labeled the cytoplasm of normal and dysplastic epithelium in both laboratories (Figure 5B-D). The dysplastic epithelium demonstrated more intense staining than morphologically normal epithelium (Figure 5D). E-cadherin, which is involved in cell-to-cell adhesion and epithelial polarity, labeled the cell membrane of normal and dysplastic epithelium (Figure 6D).

**Figure 5.**
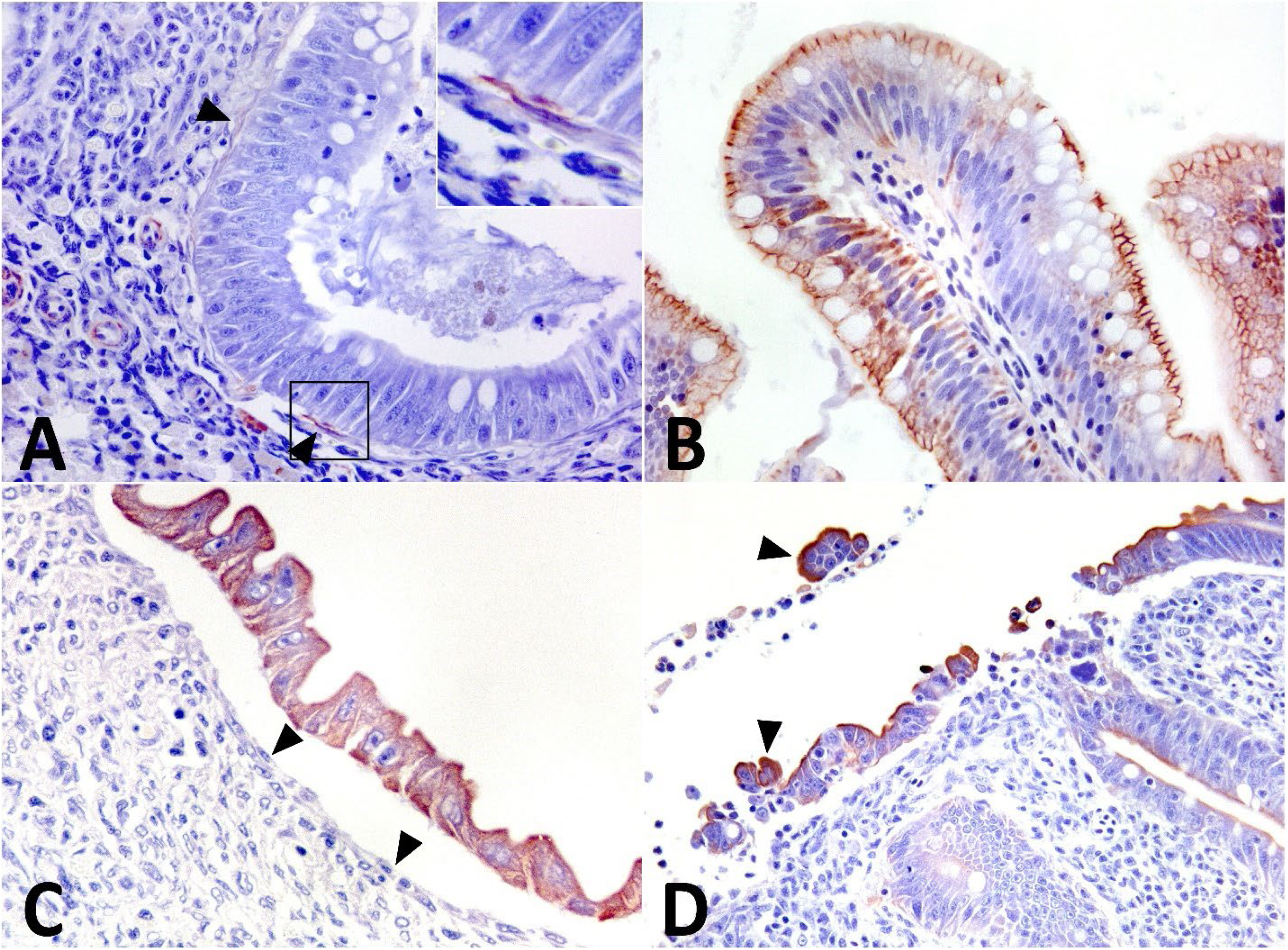
Characterization of the mucosal surface of adult salmon enteritis (ASE)-affected Chinook Salmon intestine using immunohistochemistry (IHC) for muscle actin (A) and cytokeratin WSS (B-D). A) Spindloid subepithelial cells (arrowhead) and endothelial cells of adjacent capillaries positive for muscle actin. Inset shows the spindloid subepithelial cell with positive staining for muscle actin. B) Mucosal enterocytes with positive staining for cytokeratin in healthy intestine from a non-ASE-affected broodstock. C) Mucosal enterocytes with positive staining for cytokeratin and subendothelial spindloid cells (arrowheads) with negative staining from an ASE-affected broodstock. D) Mucosal enterocytes with positive staining for cytokeratin and increased cytoplasmic staining when dysplastic or detached from the basement membrane (arrowheads) from an ASE-affected broodstock.

**Figure 6.**
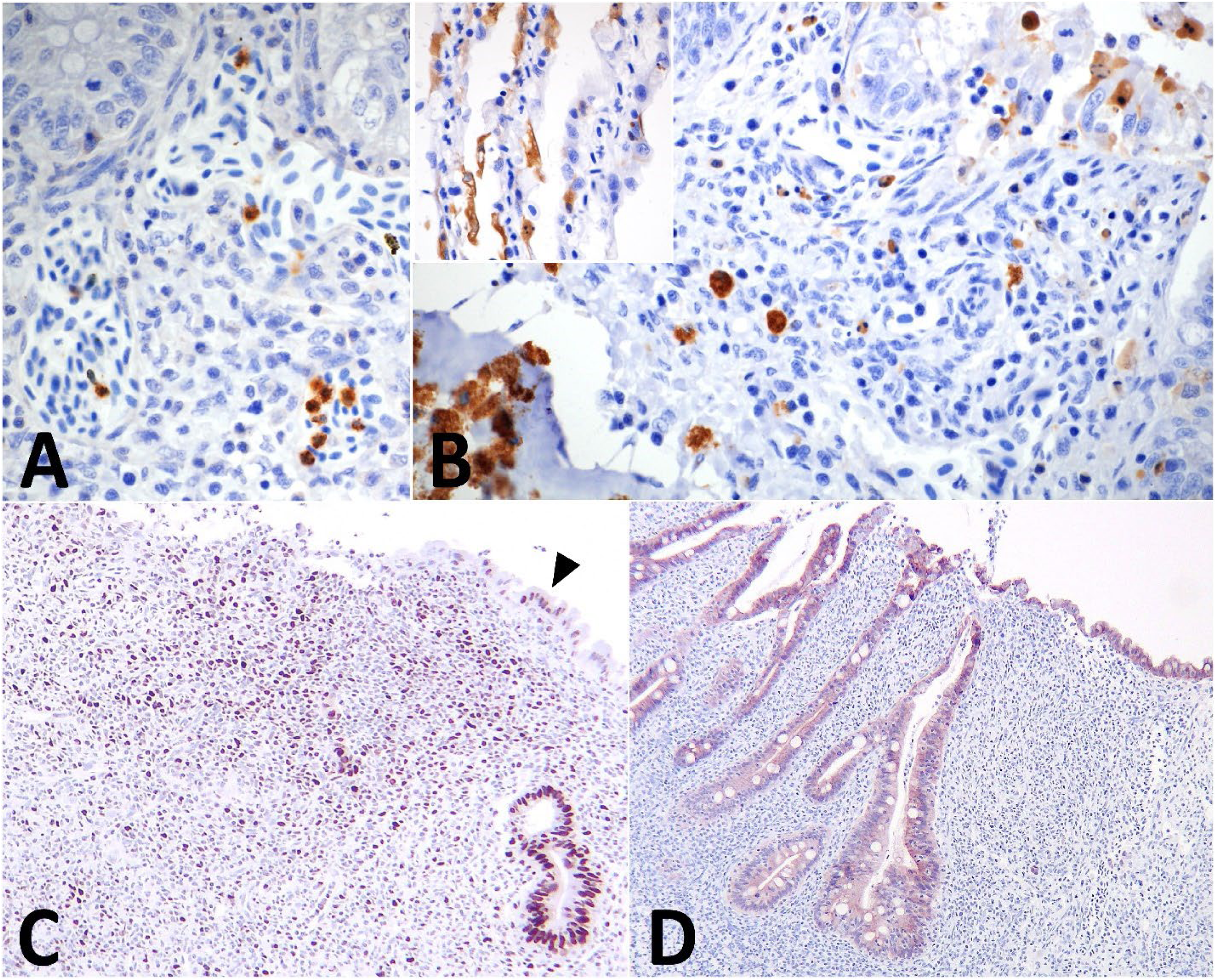
Immunohistochemistry for inflammation, apoptosis, and epithelium on adult Chinook Salmon intestine and gill with hematoxilin background stain. A) Round cells within capillaries of the lamina propria are periodically positive for TGF-beta. B) Round cells within the lamina propria, mucosa, and lumen of the intestine and granular cells within the stratum granulosum are positive for caspase 3. Within the inset, gill tissue demonstrates positive staining of gill epithelium for caspase 3. C) Nuclei of intestinal epithelium (arrowhead), round cells, and epithelium stain positive for PCNA. D) Intestinal enterocytes consistently stain positive for e-cadherin.

To further characterize the inflammatory profile of the ASE lesions, we applied IHC with antibodies for common cytokines (TGF-β, IL-1β), an apoptosis mediator (caspase 3), and cell replication factor (PCNA). The IHC for TGF-β stained individual to clustered round cells within the lamina propria (Figure 6A); no cells in the examined intestinal tissues were stained for IL-1β. Caspase 3 labelling was located in dysplastic epithelium and sloughed cells within the intestinal lumen (Figure 6B). The used antibody also labeled cytoplasmic granules of eosinophilic granular cells in the stratum granulosum, which was interpreted as non-specific cross-reactivity. The gill epithelium, which undergoes rapid turnover, had many cells stained for Caspase 3 (Figure 6 inset). The IHC for PCNA strongly labeled nuclei of cells in ASE lesions, including leukocytes, epithelium, and supporting mesenchymal cells (Figure 6C).

In more chronic lesions, the epithelium was completely ulcerated and replaced by proliferative tissue composed of fibroblasts, vascular sprouts or increased numbers of thin-walled capillaries with perpendicular orientation to the intestinal surface, mature collagenous stroma, and mixed inflammatory infiltrate as described at the beginning of this subsection (Figure 7A-B). Trichrome stained collagen light blue and revealed larger amounts of immature collagenous stroma compared to normal mucosa (Figure 7C-D), consistent with granulation tissue.

**Figure 7.**
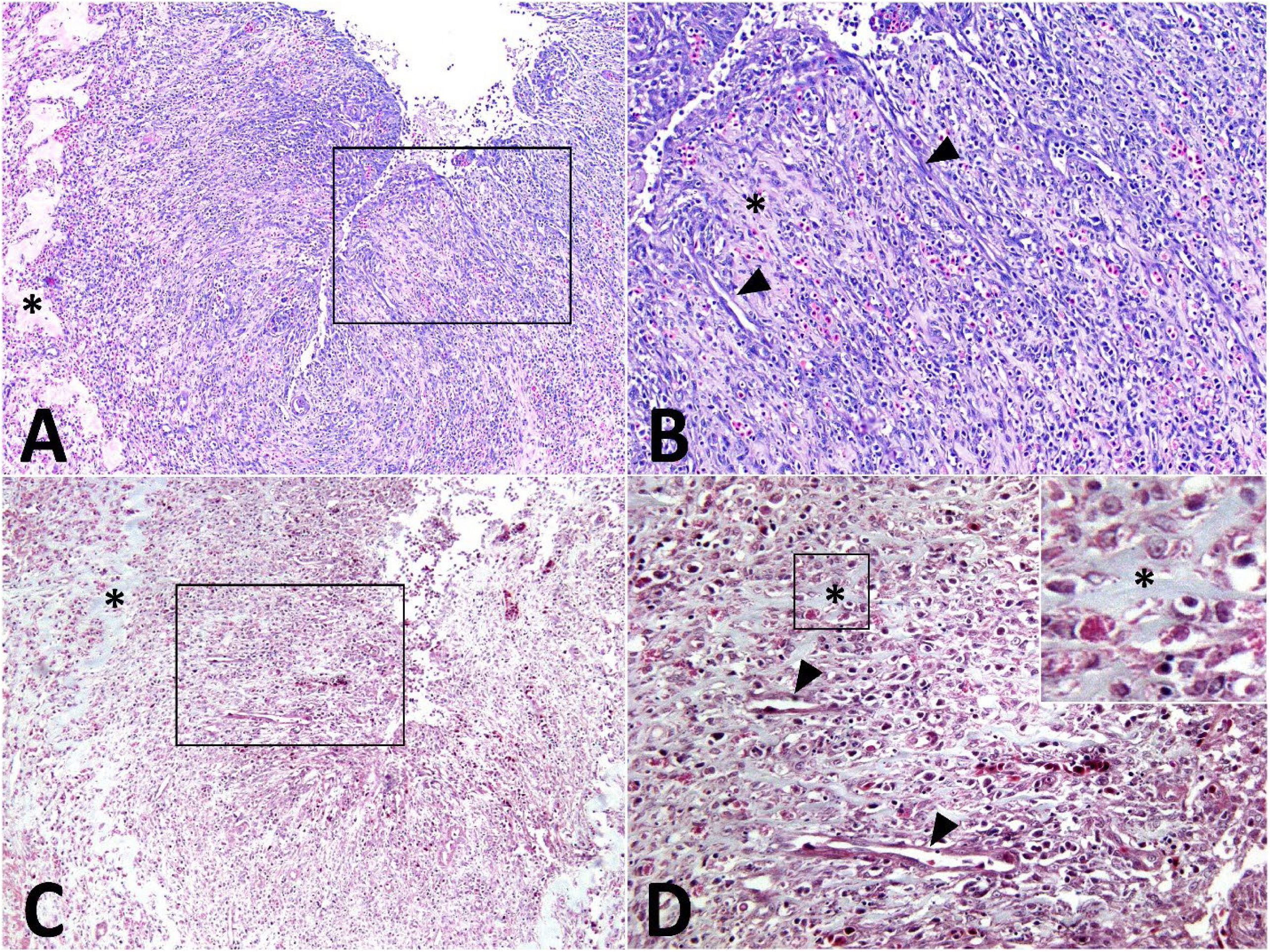
Granulation tissue in the intestine of an adult salmon enteritis (ASE)-affected Chinook Salmon. A) A severe ASE lesion showing complete ulceration of the epithelium and loss of the enteric folds with marked proliferation of tissue expanding the lamina propria and compressing the lumen. The mature collagenous stroma of the stratum compactum is indicated (*). HGE. B) Higher magnification of A (black box), showing the proliferative tissue is organized into collagenous stroma (*), reactive fibroblasts, and thin-walled capillaries (arrowheads) that are perpendicular to the ulcerated surface with a background of severe mixed inflammation. HGE. C) A sequential section of intestine from A) showing severe ASE and proliferative tissue with collagen of the stratum compactum stained blue (*). Trichrome. D) Higher magnification of C (black box), showing the same lesion as described in B) with collagen (*) stained blue and thin-walled capillaries (arrowheads) organized in a perpendicular fashion to the ulcerated surface.

#### 3.1.2 Inflammation Scores

Intestinal inflammation scores were significantly more severe in ASE-affected hatcheries compared to ASE-unaffected hatcheries (Wilcoxon rank sum test: W = 6368, p < 4.215176e-36; Table 3). ASE-affected fish had a median inflammation score of 3 (mean +/- SD: 2.85 +/- 0.41, range: 1-3), while ASE-unaffected fish had a median score of 1 (mean +/- SD: 0.57 +/- 0.55, range: 0-2). The distribution of inflammation scores differed markedly between groups (Table 3). In ASE-affected hatcheries, 87% of fish (n = 187/215) exhibited severe inflammation (score 3), 11.2% (n = 24/215) had moderate inflammation (score 2), and only 1.9% (n = 4/215) showed minimal inflammation (score 1). No ASE-affected fish had normal intestinal histology (score 0). In contrast, ASE-unaffected hatcheries showed predominantly minimal to no inflammation: 45.5% (n = 20/44) had no inflammation (score 0), 52.3% (n = 23/44) had minimal inflammation (score 1), and only 2.3% (n = 1/44) had moderate inflammation (score 2). No fish from ASE negative locations (White River and Minter Hatcheries) exhibited severe inflammation.

**Table 3.** Distribution of intestinal inflammation scores in ASE-affected and ASE-unaffected Chinook Salmon hatcheries. Inflammation was scored on a 4-point scale (0-3) based on severity and distribution of leukocytic infiltrates. Values are presented as n (%). ASE-affected populations include Carson, Round Butte, Sandy, South Santiam, and Willamette broodstock. ASE-unaffected populations include Minter Creek and White River broodstock. Asterisks (*) indicate statistically significant differences between groups for individual scores (Fisher’s exact test: p < 0.001). Overall hatchery group comparison by Wilcoxon rank sum test: W = 6368, p < 0.001.

| Hatchery Group | N | Score 0*<br>n (%) | Score 1*<br>n (%) | Score 2<br>n (%) | Score 3*<br>n (%) | Mean<br>± SD | Median | Range |
| --- | --- | --- | --- | --- | --- | --- | --- | --- |
| ASE- hatcheries | 215 | 0 (0) | 4 (1.9) | 24<br>(11.2) | 187 (87) | 2.85±<br>0.41 | 3 | 1-3 |
| Non-ASE<br>hatcheries | 44 | 20 (45.5) | 23 (52.3) | 1 (2.3) | 0 (0) | 0.57±<br>0.55 | 1 | 0-2 |

Fisher’s exact test revealed significant differences between groups with inflammation scores of 0, 1, and 3 (all p < 0.001), but not at score 2 (p = 0.091). Notably, there was complete separation at the severe inflammation level: all fish with inflammation score 3 originated from ASE-affected hatcheries, while no fish from ASE-unaffected hatcheries exhibited this level of inflammation (p < 0.001). Similarly, the absence of inflammation (score 0) was significantly associated with ASE-unaffected hatcheries (45.5% vs 0%, p < 0.001), and mild inflammation (score 1) was predominantly observed in ASE-unaffected fish (52.3% vs 1.9%, p < 0.001). Moderate inflammation (score 2) showed no significant difference between groups (11.2% vs 2.3%, p = 0.091), suggesting this represents a transitional state that may occur in both ASE-affected and ASE-unaffected populations. The total inflammation burden was substantially higher in ASE-affected hatcheries (total score: 613 across 215 fish, mean 2.85) compared to ASE-unaffected hatcheries (total score: 25 across 44 fish, mean 0.568), representing a 5-fold difference in mean cumulative inflammation severity.

Temporal variation in inflammation severity was observed among ASE-affected populations. Sandy Weir, a down-river sampling location that enables earlier-run sampling of fish migrating in the Sandy River, exhibited significantly lower inflammation scores (mean 1.8 ± 0.73, median score 2, range 1-3) compared to other ASE-affected populations (mean 2.9 ± 0.31, median score 3, range 1-3; Wilcoxon rank sum test: W = 173.5, p = 3.186119e-14). This suggests that intestinal inflammation severity may increase with disease duration or that early-stage infections present with less severe pathology.

#### 3.1.3 Lesion Distribution

In the Swiss roll arrangement (e Silva et al., 2016), ASE lesions were segmental but consistently more severe towards the orad segment, involving the pyloric ceca and first segment of the mid-intestine, with occasional extension into the second segment of the mid-intestine. The posterior segment of the intestine, in both ASE-affected and unaffected fish, routinely had mild to moderate inflammation, usually skewed more towards lymphoplasmacytic, with limited to no erosion or ulceration of the intestinal mucosa (Figure 7).

#### 3.1.4 E. schreckii and C. shasta Distribution

Neither parasite was observed in any fish from White River or Minter Creek hatcheries, which are the two populations with no evidence of ASE (Table 4). Based on the lack of parasite detection, these two hatcheries were removed from the parasite analysis presented here. *C. shasta* had an overall prevalence of 72.1% in fish from ASE-affected populations. Within the Columbia Basin, the percent epithelium present in fish with *C. shasta* (n=181) compared to fish without the parasite (n=30) was similar at 32.4% (range 0-100) vs 40.3% (range 0-100), respectively (Wilcoxon rank-sum test, p = 0.39). In contrast, *E. schreckii* was less common, affecting 25.2% (n=54/215) of total fish examined. Co-infection with both parasites occurred in 22.3% (n=48/215) of individuals. The severity of *C. shasta* infections varied considerably, with 42.8% (n=92/215) classified as mild (score 1), 16.7% (n=36/215) as moderate (score 2), and 26.5% (n=57/215) as severe (score 3). *E. schreckii* infections were predominantly mild (14.0%, n=30/215), with fewer moderate (5.6%, n=12/215) and severe (4.7%, n=10/215) cases. Some fish with severe ASE had very little if any remaining epithelium. The microsporidium has only been detected within enterocytes. Hence, with this severe epithelial loss, *E. schreckii* parasitized enterocytes were only found in sloughed cells, cells that were detached from the basement membrane, within the lumen of the intestine (Figure 8B). To further support this, fish with *E. schreckii* infection (n=54) had a mean of 53% intact epithelium (range 8-65%), compared to 27% (range 0-100%) in uninfected fish (n=161).

**Table 4.** Geographic distribution of Adult Salmon Enteritis (ASE) and enteric parasites in adult Chinook Salmon in Oregon and Washington 2022-2025 based on histologic evaluations of intestines. RDO = River Distance from the Ocean (km). n = number of fish examined. Status: S = artificially spawned at a hatchery (broodstock), P = prespawn mortality, W = collected in river at a weir. ASE % (n) = percent of sample and number of fish diagnosed with ASE. Epithelium = percent epithelium present, with mean and range. Dysplasia = mean and range dysplastic epithelium of the remaining epithelium, dysplasia was not applied if no epithelium was present. Inflam. = Intestinal inflammation score (mean and range). Average parasite score (range 0-3) and prevalence (*Ceratonova shasta* and *Enterocytozoon schreckii*) and parasite prevalence only (*Pelichonibothrium* sp. cestode larvae).

| Location | RDO<br>(km) | Month/Year | n | Status | ASE<br>% (n) | Epithelium<br>(range) | Dysplasia<br>(range) | Inflam.<br>(range) | *C. shasta* |  | *E. schreckii* |  | Cestode<br>% (n) |
| --- | --- | --- | --- | --- | --- | --- | --- | --- | --- | --- | --- | --- | --- |
|  |  |  |  |  |  |  |  |  | Score | % (n) | Score | % (n) |  |
| Willamette Basin |  |  |  |  |  |  |  |  |  |  |  |  |  |
| Willamette | 508 | 9/2022 | 15 | S | 93 (14) | 41 (0-95) | 25 (3-76) | 2.8 (2-3) | 2.2 | 93 (14) | 0.87 | 67 (10) | 33 (5) |
|  |  | 9/2023 | 25 | S | 96 (24) | 60 (10-100) | 44 (10-80) | 2.8 (2-3) | 1.5 | 92 (23) | 0.64 | 48 (12) | 20 (5) |
|  |  | 9/2024 | 10 | S | 80 (8) | 70 (41-100) | 17 (0-30) | 2.5 (2-3) | 2.2 | 100 (10) | 0.5 | 50 (5) | 40 (4) |
|  |  | 9/2025 | 20 | S | 100 (20) | 61 (0-90) | 45 (20-70) | 3.0 (3) | 0.95 | 80 (16) | 1.3 | 65 (13) | 25 (5) |
| S. Santiam | 393 | 9/2024 | 10 | S | 90 (9) | 49 (12-100) | 27 (0-60) | 2.9 (2-3) | 2.4 | 100 (10) | 0.1 | 10 (1) | 70 (7) |
| Lower Columbia Basin |  |  |  |  |  |  |  |  |  |  |  |  |  |
| Sandy | 214 | 7/2023 | 10 | W | 60 (6) | 87 (50-100) | 22 (0-80) | 1.8 (1-3) | 0.1 | 10 (1) | 0.0 | 0.0 (0) | 80 (8) |
|  |  | 9/2023 | 16 | S | 69 (11) | 61 (10-100) | 46 (0-90) | 2.6 (1-3) | 1.6 | 88 (14) | 0.38 | 19 (3) | 50 (8) |
| Carson | 278 | 9/2025 | 22 | S | 100 (22) | 28 (0-80) | 62 (20-100) | 3.0 (3) | 1.7 | 91 (20) | 0.77 | 36 (8) | 41 (9) |
| Middle Columbia Basin |  |  |  |  |  |  |  |  |  |  |  |  |  |
| Round Butte | 469 | 8/2022 | 11 | P | 100 (11) | 3.0 (0-20) | 78 (10-100) | 3.0 (3) | 0.91 | 64 (7) | 0.0 | 0.0 (0) | 0.0 (0) |
|  |  | 9/2022 | 19 | S | 100 (19) | 7.0 (0-60) | 76 (0-100) | 3.0 (3) | 2.4 | 100 (19) | 0.0 | 0.0 (0) | 0.0 (0) |
|  |  | 6-8/2023 | 13 | P | 100 (10) | 1.2 (0-10) | 90 (80-100) | 3.0 (3) | 2.4 | 100 (13) | 0.1 | 7.7 (1) | 15 (2) |
|  |  | 8/2023 | 14 | S | 100 (14) | 1.9 (0-15) | 75 (50-100) | 3.0 (3) | 1.2 | 100 (14) | 0.0 | 0.0 (0) | 7.1 (1) |
|  |  | 7-8/2024 | 10 | P | 100 (10) | 1.0 (0-13) | 10 (0-20) | 3.0 (3) | 1.5 | 100 (10) | 0.0 | 0.0 (0) | 20 (2) |
|  |  | 8/2025 | 20 | S | 100 (20) | 3.1 (0-37) | 73 (50-100) | 3.0 (3) | 0.8 | 70 (14) | 0.1 | 5.0 (1) | 0.0 (0) |
| Puget Sound |  |  |  |  |  |  |  |  |  |  |  |  |  |
| White River | 201 | 9/2022 | 15 | S | 0.0 (0) | 100 (100) | 0.0 (0) | 0.67 (0-2) | 0.0 | 0.0 (0) | 0.0 | 0.0 (0) | 73 (11) |
|  |  | 9/2023 | 10 | S | 0.0 (0) | 100 (100) | 0.0 (0) | 1.0 (1) | 0.0 | 0.0 (0) | 0.0 | 0.0 (0) | 90 (9) |
| Minter Creek | 2 | 9/2023 | 19 | S | 0.0 (0) | 100 (100) | 0.0 (0) | 0.26 (0-1) | 0.0 | 0.0 (0) | 0.0 | 0.0 (0) | 74 (14) |

**Figure 8.**
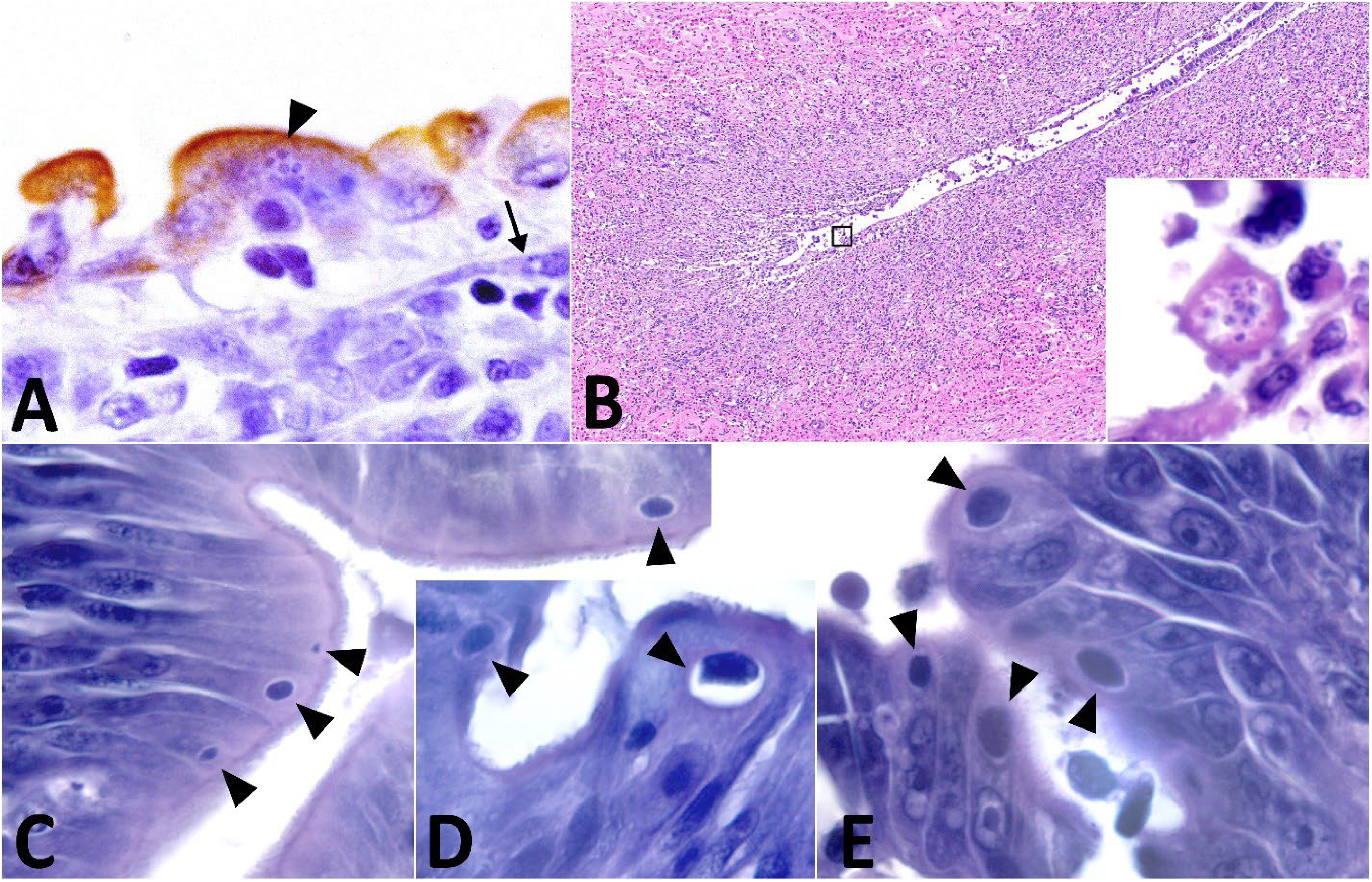
*Enterocytozoon schreckii* in enterocytes from Chinook Salmon intestines affected by adult salmon enteritis (ASE). A) Willamette Hatchery broodstock. Mature spores within the cytoplasm of an enterocyte (arrowhead), which positively stains for cytokeratin WSS and a subepithelial myofibroblast that fails to stain for WSS (arrow). WSS and hematoxylin. B) Round Butte Hatchery broodstock. Severe ASE lesion with marked ulceration of the intestine and mixed inflammation. Inset displays an enterocyte detached from the epithelial basement membrane containing mature spores. HGE. C-E) Presporogonic stages (arrowheads) from C) Willamette Hatchery broodstock, D) Carson Hatchery, and E) Round Butte Hatchery. HGE.

**Figure 9.**
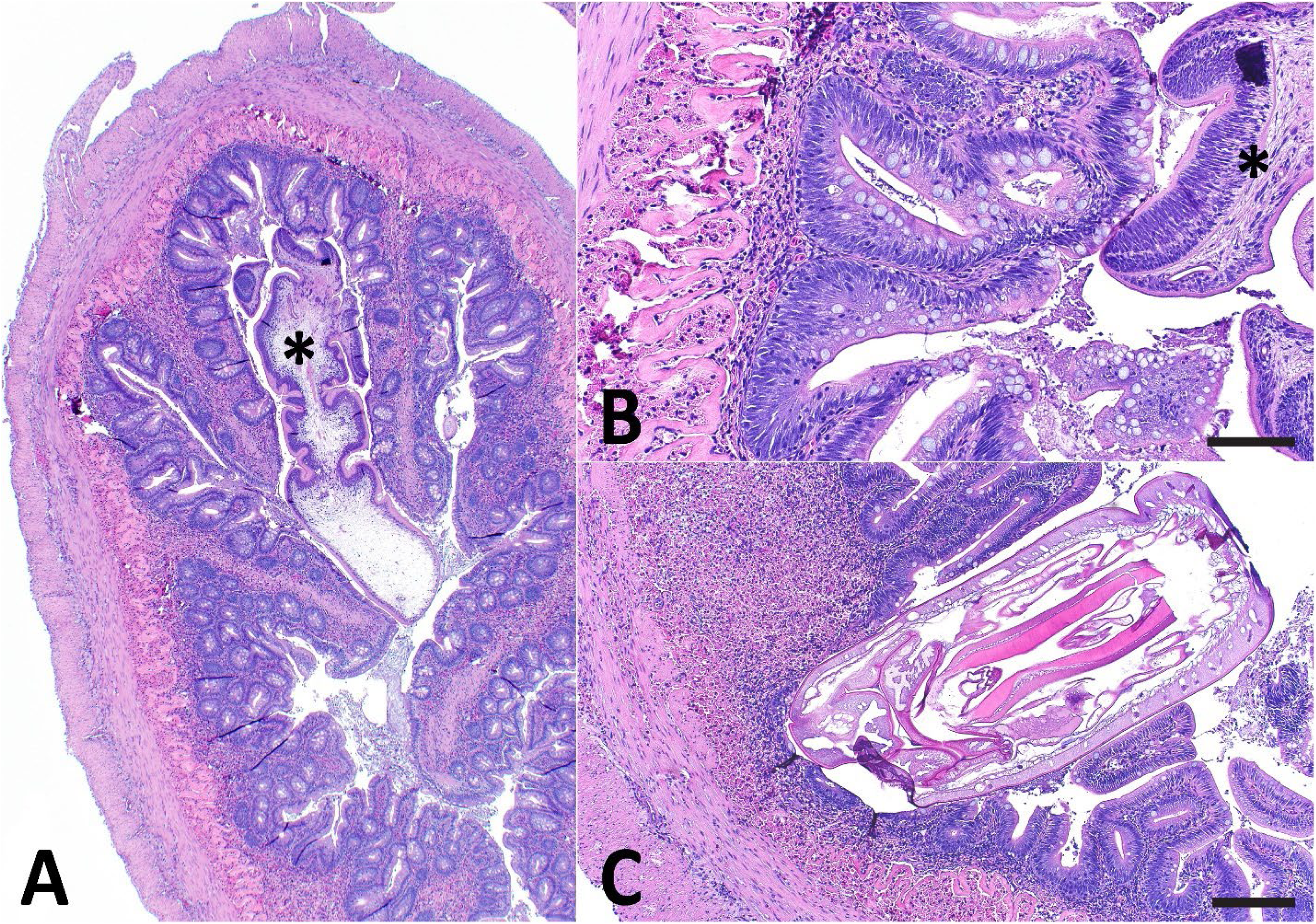
Macroparasites found in the intestines of Chinook Salmon broodstock. (A) Cestode (*) within the intestinal lumen of a White River fish, and at higher power (B) showing the attachment point of the cestode (*) to the mucosal surface with a mild enteritis (score 1). Scale bar = 100 um. (C) An acanthocephalan penetrating the mucosa and lamina propria of the intestine of a Minter Creek fish with a localized granulocytic inflammatory response. Scale bar = 200 um. HGE.

Parasite prevalence varied markedly among hatcheries, from absent in those without ASE to >80% in the those with ASE (Table 4). *C. shasta* prevalence was highest at Santiam (100%, n=10/10), Carson (91%, n=20/22), Willamette (90%, n=63/70), Round Butte (86%, n=77/87), and Sandy Hatchery (88%, n=14/16). In contrast, Minter Creek and White River showed no *C. shasta* infections (0%, n=0/19 and n=0/25, respectively), while Sandy Weir had minimal prevalence (10.0%, n=1/10). *E. schreckii* showed a different distribution pattern, with highest prevalence at Willamette (57%, n=40/70), Carson (36%, n=8/22), and Sandy Hatchery (19%, n=3/16).

The presence of both parasites was strongly associated with intestinal inflammation. Fish with no inflammation (score 0) or minimal inflammation (score 1) showed no parasite infections. Fish with moderate inflammation (score 2) had intermediate prevalence: 72.0% (n=18/25) for *C. shasta* and 24.0% (n=6/25) for *E. schreckii*. Among fish with severe inflammation (score 3), *C. shasta* prevalence was 89.3% (n=167/187) and *E. schreckii* prevalence was 24.6% (n=46/187).

In contrast, *E. schreckii* infections had a distinct anatomical distribution within the intestinal tract. The parasite was predominantly identified in the pyloric ceca and mid-intestine regions, where characteristic presporogonic stages and spores were visible within the cytoplasm of epithelial cells or enterocytes. The localization of the spores in epithelial cells was confirmed using a pancytokeratin IHC (Figure 8A). In contrast, *E. schreckii* was notably absent in the posterior segment. This pyloric ceca and mid intestinal localization pattern was consistent across infected individuals, regardless of infection severity.

### 3.2 Geographic Distribution

The Willamette and Columbia River populations all had a high prevalence of ASE in their broodstock, as well as *C. shasta* and *E. schreckii* (Table 4). In contrast, the two rivers flowing into Puget Sound had no ASE or these two parasites. Larval cestodes were noted in the lumen of the intestine in several fish from both positive and negative ASE locations (Figure 6). Based on histologic sections, the worm had four suckers. Our previous study (Polley et al., 2026), with examination of whole worms in wet mounts, identified this worm as a member of the genus *Pelichnibothrium*.

The following summarizes intestinal changes and pathogens based on specific watersheds. We also provide a description of each sampling location with data on historic PSM levels provided by hatchery staff and other references. First described are Puget Sound populations, White River and Minter Creek fish, as they represent populations without ASE. Then the Columbia River Basin populations, including the other extreme for ASE, Round Butte, with consistently the most severe ASE based on destruction of the intestinal epithelium. ASE-affected systems presented changes more consistent with our earlier description of ASE, along with *C. shasta* and *E. schreckii* infections (Nervino et al., 2024).

#### 3.2.1 Puget Sound Populations

The two Puget Sound populations (White River Hatchery, n=25; and Minter Creek Hatchery, n=16) represent geographically distant Chinook Salmon populations that are not connected to the Columbia River basin. White River Hatchery is located below the Mud Mountain Dam on the White River, which is a tributary of the Puyallup River. It is located 33 river miles upstream from Commencement Bay in the Puget Sound. Spring Chinook Salmon begin returning to the hatchery in May, and spawning at the hatchery begins in August and ends in October. Broodstock are held on a mix of river and well water prior to spawning. Typical PSM at this facility is under 10% for the entire holding period; historically the majority of PSM has been caused by saprolegniasis and furunculosis, and fish are treated with formalin (167 ppm for 1 hour daily to weekly) and occasionally antibiotic injections. The summer water temperatures range from about 13-18 °C.

Minter Creek Hatchery is located one river mile up Minter Creek, which connects directly to Henderson Bay in the Puget Sound. Spring Chinook Salmon begin returning to the hatchery in June, and spawning at the hatchery begins in September and ends by October. Typical PSM at this facility is low; fish are treated with formalin and salt to control PSM. Here fish are held in tanks at 10-15°C. PSM at this facility is only occasionally associated with outbreaks of *Ichthyophthirius multifillis*. Both of these populations exhibited markedly different pathogen profiles compared to Columbia River populations, with no detection of ASE, *C. shasta*, or *E. schreckii* in any of the total 44 fish examined (Table 4).

An interesting finding by histopathology in three Minter Creek fish was a profound proliferation of eosinophilic granular cells (EGCs) or granulocytes expanding and infiltrating all layers of the intestinal tissues including the mucosa, submucosa, muscularis, and serosa, described as severe diffuse transmural eosinophilic granulocytic enteritis (Figure 10). A similar unusual eosinophilic granule cell proliferation was described by Kent et al. (Kent et al., 1663) in coho salmon, with some theorized causes including neoplasia, oncogenic viruses, and endoparasitism.

**Figure 10.**
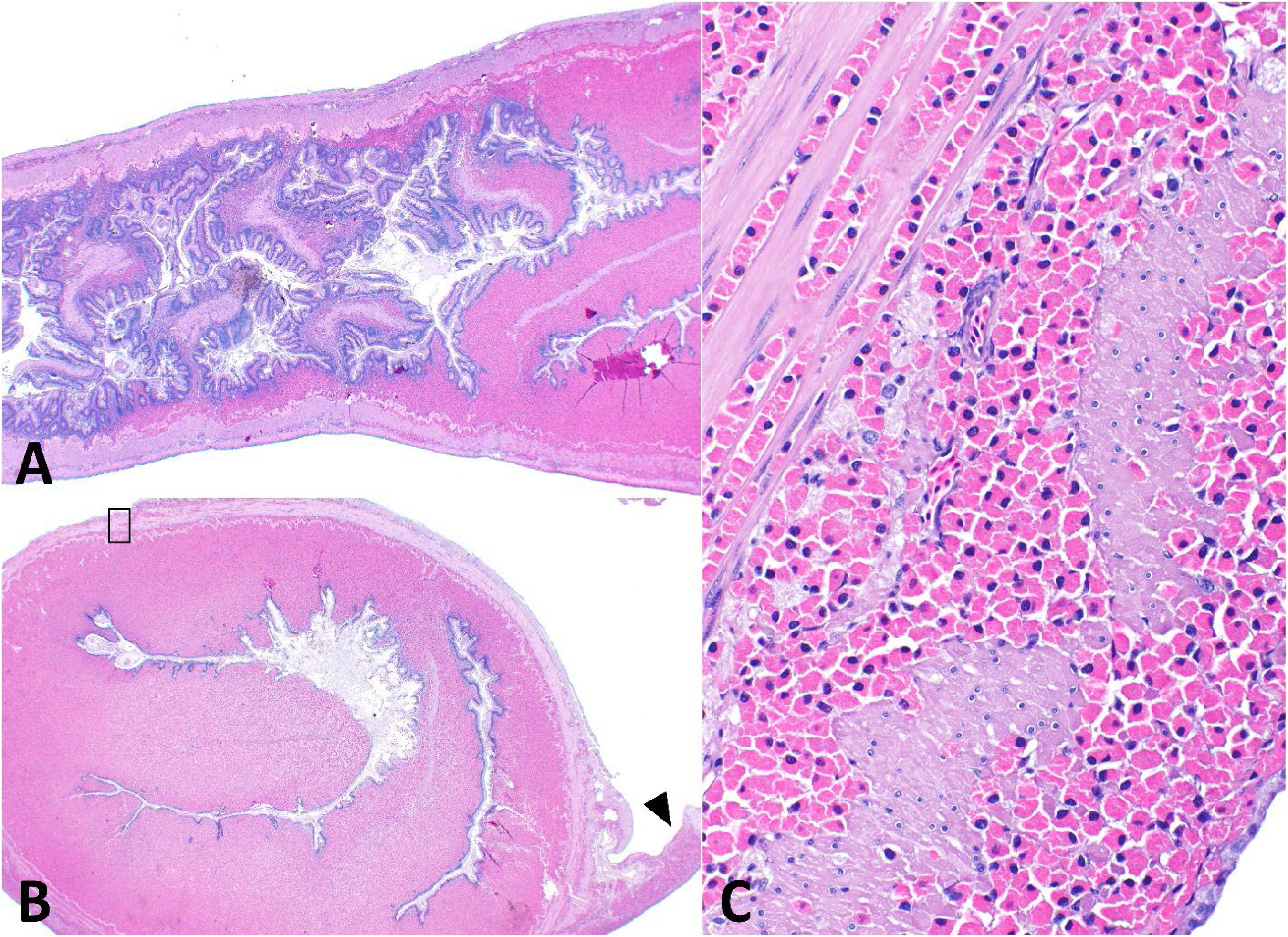
Eosinophilic granular cell (ECG) or granulocytic proliferative lesion within the mid-intestine of a Minter Creek Chinook Salmon. (A) Sagittal and (B) transverse section of mid-intestine with severe expansion of the lamina propria by a round cell population, which extends into the associated mesentery (arrowhead). Higher power of this lesion is shown in (C) and indicated by the black box. The round cells contain brightly eosinophilic cytoplasmic granules, a single round eccentric nucleus with moderately expanded chromatin, and frequent singular nucleoli. The infiltrative population extends to and expands the inner circular and out longitudinal muscularis with cells periodically seen within the serosal layer.

Other helminths were also seen by histopathology while examining fish from the Puget Sound. One fish from Minter Creek Hatchery showed the anterior portion of an acanthocephalan embedded within the mucosal wall (Figure 6C). Several fish from White River had larval nematodes embedded within the submucosa or muscular wall of the intestine surrounded by well-defined granulomas, sometimes with peritonitis. Larval cestodes were highly prevalent in both Puget Sound populations (80% in White River, n=20/25; 73.68% in Minter Creek, n=14/16). Despite high cestode burdens, intestinal architecture remained intact, with no evidence of the severe epithelial damage observed in Columbia Basin populations.

The Puget Sound populations demonstrated a distinct pathogen profile characterized by the absence of ASE, *C. shasta*, and *E. schreckii* (Table 4; Figure 2). This contrasts sharply with Columbia River Basin populations and suggests geographic isolation, at least based on histology, from the primary transmission sources for these pathogens in Oregon.

#### 3.2.2 Columbia River Basin Populations

The Columbia River basin populations exhibited high prevalence of ASE and the associated parasites, *C. shasta* and *E. schreckii*, in stark contrast to the Puget Sound populations. These sampling locations are organized by watershed (Table 1), including the hatcheries from Lower Columbia Basin (Carson and Sandy), the Middle Columbia Basin (Round Butte), and the Willamette Basin (South Santiam and Willamette).

Carson Hatchery is located at river mile 18 on the Wind River in the Lower Columbia Basin watershed (Figure 1), which enters the Columbia River 155 miles upstream from the Pacific Ocean. River water temperatures in the summer are high, and hence the adult salmon are held in water derived from a spring at 8-10 °C. PSM has been low at this facility in recent years. We examined 22 broodstock fish (Table 4), which did not exhibit obvious macroscopic internal changes. Histology revealed consistent severe intestinal inflammation (mean score 3.00) and loss of the epithelium across all fish examined. The mean percent epithelium present was 27.9%, indicating substantial epithelial loss, with mean dysplasia of 61.6% in the remaining epithelium. Hence, all the fish were diagnosed as positive for ASE. *C. shasta* was highly prevalent (91.0%). *E. schreckii* was detected, based on presence of spores in 8 fish (36.4%). Enterocytes with varying sizes of presporogonic forms were generally more numerous than those containing spores (Figure 8C-E). Larval cestodes were observed in 6 fish (40.6%).

The Sandy River flows directly into the Columbia River within the Lower Columbia Basin watershed. In 2023, adults returning to the Sandy Hatchery were captured early in the summer, transported by trucks to the Clackamas Hatchery. It is located on the Clackamas River, which flows into the Willamette River system at river mile 24.8 (Figure 1). Summer water temperatures at the hatchery range 13-18 °C. Limited data are available on PSM in the Sandy River, but it has historically been low. Sandy River broodstock fish (n=16) (Table 4) exhibited severe inflammation (mean score 2.62), with ASE diagnosed in 75% of the fish. *Ceratonova shasta* was detected in 87.5% (n=14/16) of fish and *E. schreckii* in 18.7% (n=3/16). Cestodes were present in 50.0% (n=8/16) of fish. Mean epithelium remaining was 60.6% with mean dysplasia of 45.80%. The Sandy Weir (n=10), a sampling location on the Bull Run River located earlier in the spawning migration on the Sandy River, allowed for collection of adult salmon in July 2023, which is two months earlier than natural spawning. Mean inflammation score was 1.80, with 87.0% epithelium remaining and 22.0% dysplasia, representing the mildest pathologic changes among Columbia River Basin populations. Hence, based on our designated cut-offs, 30% were diagnosed with ASE. Nevertheless, linking observations with those in the same population at spawning at the Clackamas Hatchery, most of the fish could be diagnosed as an early form of ASE. These earlier run fish showed a distinct pattern, with lower *C. shasta* prevalence (10.0%) and no *E. schreckii* detection. However, cestode prevalence was high, in 8 of 10 fish.

Round Butte Hatchery is located on the Deschutes River in the Middle Columbia Basin watershed (Figure 1). This hatchery has a spring fed water source and summer water temperatures are consistently around 10°C for holding spring Chinook Salmon broodstock. Adults are collected from the Deschutes River from May through August, and fish are spawned in mid-August-September. PSM in adult Chinook Salmon held at the hatchery was quite variable over the years of this study, with a decline over the years: 2022, 54%; 2023, 16%; 2024, 13%; and 2025, 4.8%. PSM and post-spawned broodstock were collected from 2022 to 2025 (n=87) (Table 4). Round Butte Hatchery demonstrated the most severe intestinal pathologic changes among all populations examined. Mean inflammation score was 3.00, with extensive epithelial destruction evident by a mean of only 3.2% epithelium remaining. The remaining epithelium showed severe dysplasia (mean 52.3%).

*Ceratonova shasta* prevalence was 88.5% of the fish (Table 4). Whereas severe ASE occurred across all samples, there was a variable prevalence and severity scores for *C. shasta* across the years, as demonstrated with individual fish scores in the raw data (https://doi.org/10.7267/gt54kx16f). The infection, based on intestinal histology scores, was less prevalent and severe in 2025, reduced from 100% to 75%, and most of the infected fish had lighter infections (score 1). Whereas not a primary focus of this study, it was noted that many fish at Round Butte Hatchery had extraintestinal *C. shasta*.

With minimal epithelium in the Round Butte broodstock, only two fish from 2023 had clearly detectable *E. schreckii* infections based on the observation of mature spores within detached, luminal enterocytes (Figure 8B). One fish from 2025, Fish 8, which had some epithelium present, exhibited subspherical structures in the apical aspect of epithelial cells consistent with presporognic stages of *E. schreckii* (Figure 8E). This included one in which a partial halo was seen due to the early-stage parasites withdrawn from the cyst wall. This is a processing artifact described previously (Nervino, 2022; Nervino et al., 2024). Cestodes were rare, detected in only 5 fish.

While performing necropsies at Round Butte Hatchery, we noted that both PSM and post-spawned broodstock exhibited reno-splenomegaly, sometimes with mottled pallor, which we recently described in Polley et al. (2026) as due to extraintestinal *C. shasta* infections. The independent clinical examinations of PSM broodstock conducted by one of us (S. S.) provided additional descriptive data. Here infectious status for *C. shasta*, based on wet mounts for myxospores from kidney or intestine, and *R. salmoninarum*, by direct fluorescence antibody testing (DFAT), were available from 2022-2025 for 45 PSM broodstock. Extra-intestinal infections with *C. shasta* were confirmed in 29% (n=13/45) of the PSM broodstock, with mottled discoloration documented in 2 fish, both from 2023 with myxospores present in the kidney wet mount, and friable kidneys documented in 5 fish from 2022. Macroscopic renal changes were seen in 13% of the fish (n=6/45), but myxospores were not detected in two of these fish. Two fish were diagnosed with *R. salmoninarum* infections. We had corresponding histology of intestines on 16 of these PSM broodstock (https://doi.org/10.7267/gt54kx16f). *C. shasta* was diagnosed by histology in the intestine of all these fish (n=16/16) and by wet mounts for myxospores in the kidney of 11 fish. In 2025, we observed that essentially all broodstock fish that we necropsied had renomegaly, which we presume was caused by *C. shasta* because *R. salmoninarum* ELISA tests were negative. However, histology of the intestine showed that many were negative for the parasite or had a score of 1. A total of 11 fish were available to compare wet mounts to histology of intestines from the same fish. Whereas all were positive by histology, five were negative by wet mount.

The South Santiam River flows into the Willamette River near Salem, Oregon with the Willamette Basin watershed (Figure 1), with summer water temperature ranging from about 15-21°C. PSM in this river historically has been high, up to 72%, with a mean of 22% for 2005-2017 (Bowerman et al., 2018). In 2024, we examined 10 fish from South Santiam Hatchery (Table 4), and they presented patterns of lesions diagnostic for ASE, as seen in other stocks of fish from the Willamette system. Mean inflammation score was 2.90, with 48.5% epithelium remaining and 26.50% dysplasia. One fish was diagnosed as negative for ASE with 100% intact epithelium, no epithelial dysplasia, and an intestinal inflammation score of 2. All fish were infected with *C. shasta*, while *E. schreckii* was detected in only one fish. Cestodes were present in 70.0%.

Willamette Hatchery is located along Salmon Creek, which is three miles upstream of its confluence with the Middle Fork of the Willamette River within the Willamette Basin watershed (Figure 1). PSM has historically been very high in the Willamette River, with a mean of 80% from 2006-2017 (Bowerman et al., 2018). Summer water temperatures here range from 15-21 °C. In the past, this population spawned at the Willamette Hatchery, and these represent our samples from 2022-2023. However, for samples collected in 2025, broodstock were trapped at Dexter Reservoir, located 16 miles southeast of Eugene, Oregon on the Middle Fork Willamette River. Broodstock were then transported and held at the South Santiam Hatchery before spawning. Over the four years of sampling, fish from this river consistently exhibited ASE in ≥90% of the broodstock according to our statistical cutoff. Across all broodstock (n=70), the mean inflammation score was 2.81 (range 2-3), with 57.6% epithelium remaining (range 0-100%) and 36.25% dysplasia. They also showed high *C. shasta* prevalence (90.0%, n=63/70) and the highest *E. schreckii* prevalence among all populations (56.52%). Also, early *E. schreckii* meront stages were far more numerous than mature spores, suggesting earlier development, which also was repeated in the Carson Hatchery. Cestodes were detected in 27.1% of the broodstock.

#### 3.2.3 Geographic Patterns and Synthesis

The geographic distribution of ASE and associated parasites revealed a clear biogeographic divide between Puget Sound and Columbia River Basin populations. *Ceratonova shasta* and *E. schreckii* were highly prevalent in most Columbia Basin populations, with 85.6% for *C. shasta* and 23.4% for *E. schreckii*. Concurrently, the fish exhibited severe intestinal changes consistent with ASE, characterized by inflammation, epithelial loss, and dysplasia. In contrast, Puget Sound populations (n=44) showed complete absence of ASE, *C. shasta*, and *E. schreckii* (0%, n=0/44 for all three conditions). This parasitological divide corresponded to dramatic differences in intestinal pathology. Puget Sound fish (n=44) showed minimal inflammation (mean 0.26-0.8), complete epithelial retention (mean 100%), and no dysplasia (mean 0%). In contrast, most Columbia Basin fish (n=222) showed severe inflammation (mean 2.62-3.0), extensive epithelial loss (mean retention 3.2-87%), and widespread dysplasia (mean 26.5-61.58%).

Within the Columbia Basin, *E. schreckii* prevalence showed substantial variation among hatcheries based on histologic detection. Willamette fish showed the highest prevalence (55.71%), followed by Carson (36.36%), Sandy Hatchery (18.75%), Santiam (10%), Round Butte (1.15%), and Sandy Weir (0.00%). Variation in intestinal lesion severity was also evident within the Columbia Basin hatcheries. Round Butte consistently showed the most severe epithelial erosion and ulceration (mean 3.2% epithelium remaining), while the Sandy Weir exhibited the mildest pathology (mean 87.0% epithelium remaining). This variation may reflect differences in pathogen exposure, disease progression, or host factors among populations, despite their shared connection to the Columbia River system.

Cestode prevalence demonstrated an inverse geographic pattern, with significantly higher prevalence in Puget Sound (72.7%, n=32/44) compared to Columbia River Basin populations (23.0%, n=51/222). Notably, high cestode prevalence in Puget Sound populations was not associated with severe pathologic changes, suggesting cestodes alone do not drive the epithelial damage characteristic of ASE.

## 4 Discussion

### 4.1 ASE Description and Pathology

This study builds upon the initial characterization of adult salmon enteritis (ASE) by Nervino et al. (2024), providing a more comprehensive pathological description through expanded histological techniques and geographic sampling. Using histochemical stains and IHC, we further defined the progression, intestinal localization, and inflammatory profile of ASE lesions. Antibodies developed against mammalian tissues and cellular targets, and not specifically intended for fish, are now widely used in teleosts owing to cross-reactivity. Hence, their performance varies, with some showing acceptable reactivity and others failing to do so (Antuofermo et al., 2023). Table 5 provides a review of selected IHCs and references pertinent to this study.

**Table 5.**
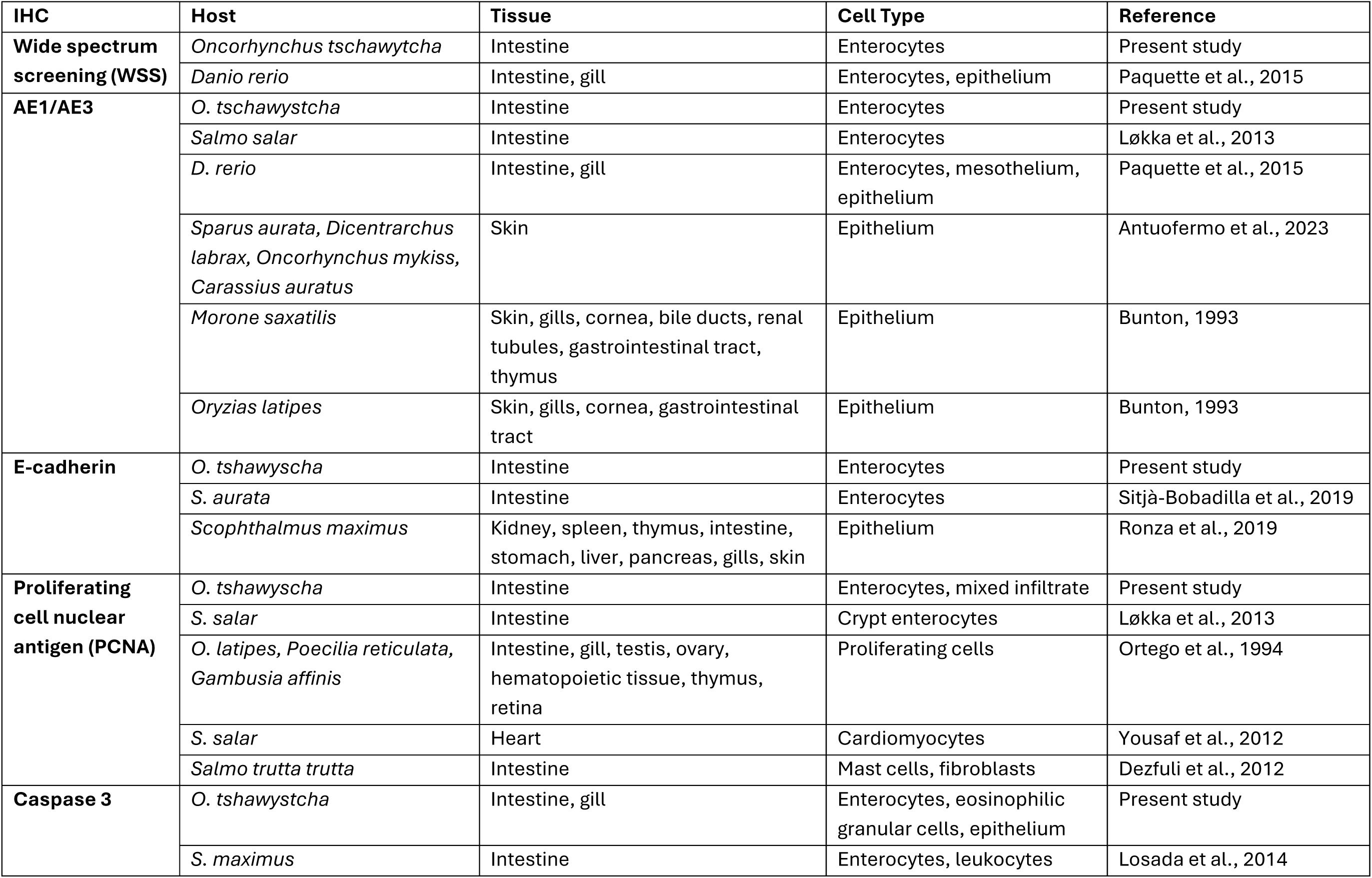
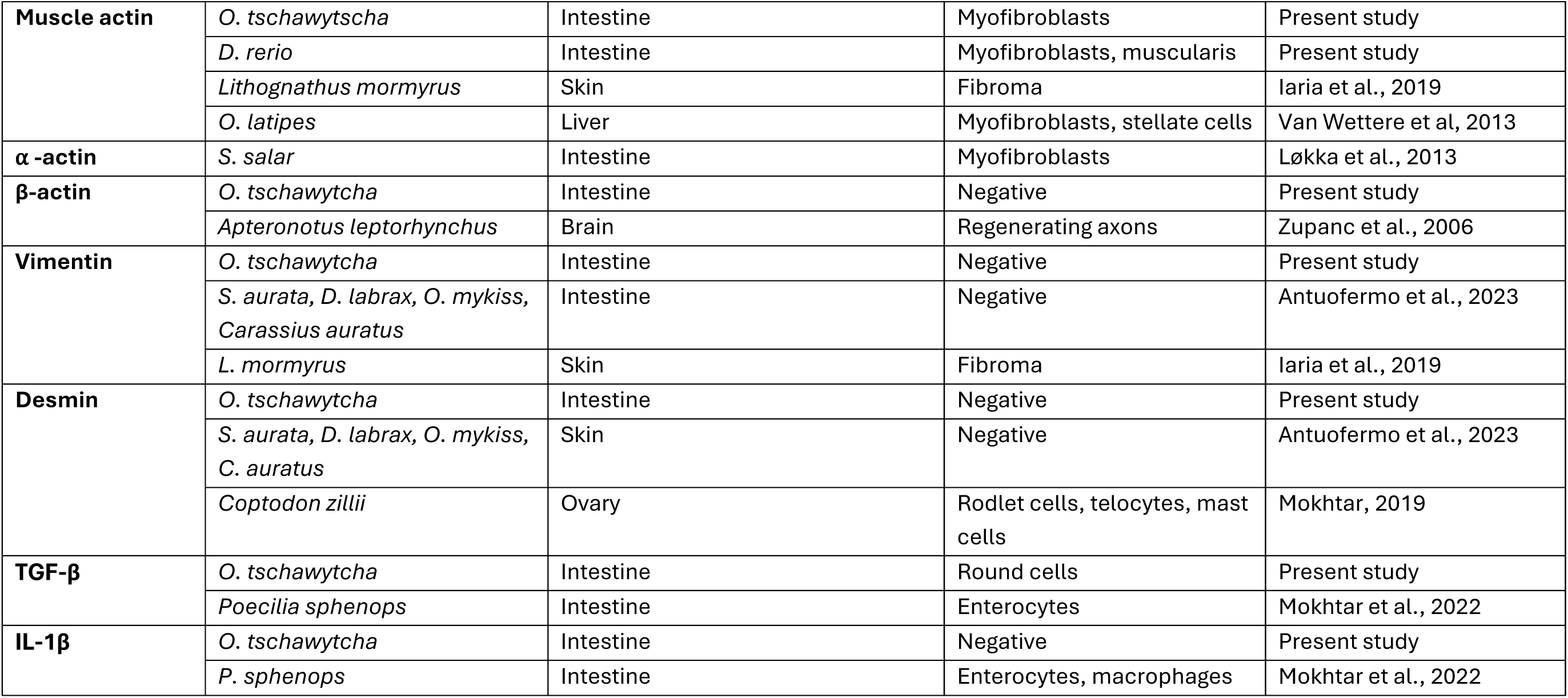
Selected review of IHCs used with teleost fishes in light microscopy pertinent to the present investigation of Adult Salmon Enteritis in Chinook Salmon (*Oncorhynchus tshawytscha)*.

| IHC | Host | Tissue | Cell Type | Reference |
| --- | --- | --- | --- | --- |
| **Wide spectrum screening (WSS)** | *Oncorhynchus tshawytscha* | Intestine | Enterocytes | Present study |
|  | *Danio rerio* | Intestine, gill | Enterocytes, epithelium | Paquette et al., 2015 |
| **AE1/AE3** | *O. tshawytscha* | Intestine | Enterocytes | Present study |
|  | *Salmo salar* | Intestine | Enterocytes | Løkka et al., 2013 |
|  | *D. rerio* | Intestine, gill | Enterocytes, mesothelium, epithelium | Paquette et al., 2015 |
|  | *Sparus aurata*, *Dicentrarchus labrax*, *Oncorhynchus mykiss*, *Carassius auratus* | Skin | Epithelium | Antuofermo et al., 2023 |
|  | *Morone saxatilis* | Skin, gills, cornea, bile ducts, renal tubules, gastrointestinal tract, thymus | Epithelium | Bunton, 1993 |
|  | *Oryzias latipes* | Skin, gills, cornea, gastrointestinal tract | Epithelium | Bunton, 1993 |
| **E-cadherin** | *O. tshawytscha* | Intestine | Enterocytes | Present study |
|  | *S. aurata* | Intestine | Enterocytes | Sitjà-Bobadilla et al., 2019 |
|  | *Scophthalmus maximus* | Kidney, spleen, thymus, intestine, stomach, liver, pancreas, gills, skin | Epithelium | Ronza et al., 2019 |
| **Proliferating cell nuclear antigen (PCNA)** | *O. tshawytscha* | Intestine | Enterocytes, mixed infiltrate | Present study |
|  | *S. salar* | Intestine | Crypt enterocytes | Løkka et al., 2013 |
|  | *O. latipes*, *Poecilia reticulata*, *Gambusia affinis* | Intestine, gill, testis, ovary, hematopoietic tissue, thymus, retina | Proliferating cells | Ortego et al., 1994 |
|  | *S. salar* | Heart | Cardiomyocytes | Yousaf et al., 2012 |
|  | *Salmo trutta trutta* | Intestine | Mast cells, fibroblasts | Dezfuli et al., 2012 |
| **Caspase 3** | *O. tshawytscha* | Intestine, gill | Enterocytes, eosinophilic granular cells, epithelium | Present study |
|  | *S. maximus* | Intestine | Enterocytes, leukocytes | Losada et al., 2014 |
| **Muscle actin** | *O. tschawytsha* | Intestine | Myofibroblasts | Present study |
|  | *D. rerio* | Intestine | Myofibroblasts, muscularis | Present study |
|  | *Lithognathus mormyrus* | Skin | Fibroma | Iaria et al., 2019 |
|  | *O. latipes* | Liver | Myofibroblasts, stellate cells | Van Wettere et al, 2013 |
| **α-actin** | *S. salar* | Intestine | Myofibroblasts | Løkka et al., 2013 |
| **β-actin** | *O. tschawytcha* | Intestine | Negative | Present study |
|  | *Apteronotus leptorhynchus* | Brain | Regenerating axons | Zupanc et al., 2006 |
| **Vimentin** | *O. tschawytcha* | Intestine | Negative | Present study |
|  | *S. aurata*, *D. labrax*, *O. mykiss*,<br>*Carassius auratus* | Intestine | Negative | Antuofermo et al., 2023 |
|  | *L. mormyrus* | Skin | Fibroma | Iaria et al., 2019 |
| **Desmin** | *O. tschawytcha* | Intestine | Negative | Present study |
|  | *S. aurata*, *D. labrax*, *O. mykiss*,<br>*C. auratus* | Skin | Negative | Antuofermo et al., 2023 |
|  | *Coptodon zillii* | Ovary | Rodlet cells, telocytes, mast cells | Mokhtar, 2019 |
| **TGF-β** | *O. tschawytcha* | Intestine | Round cells | Present study |
|  | *Poecilia sphenops* | Intestine | Enterocytes | Mokhtar et al., 2022 |
| **IL-1β** | *O. tschawytcha* | Intestine | Negative | Present study |
|  | *P. sphenops* | Intestine | Enterocytes, macrophages | Mokhtar et al., 2022 |

The consistent microanatomic findings of ASE include severe mixed inflammation, including granulocytes, and progressive intestinal mucosal damage leading to epithelial erosion and ulceration, with attempted repair through proliferative granulation tissue. In regions of ulceration, the surface is covered by elongate, spindloid cells we identified as myofibroblasts through muscle actin immunoreactivity and absence of staining with epithelial cell markers (WSS or e-cadherin). These putative myofibroblasts were consistent in morphology and location with those described in Atlantic Salmon by Løkka et al. (2013). Attenuated enterocytes, perhaps attempting to cover ulcerated regions, could be confused with myofibroblasts. However, normal and dysplastic epithelial cells stained readily with WSS (Figure 5D; Figure S1) and e-cadherin. Indeed, normal enterocytes showed staining more concentrated in the apical cytoplasm, whereas degenerate, dysplastic, detached, or otherwise abnormal enterocytes consistently displayed more intense staining throughout the cytoplasm. There are several reports of IHCs used to identify epithelial cells in teleost fish (Table 5), and in our study WSS showed stronger specific reactivity than AE1/AE3 for intestinal mucosa of Chinook Salmon.

Examination of intestinal Swiss roll sections revealed that ASE lesions are not uniformly distributed throughout the intestinal tract. Lesions were most severe in the mid-intestine and pyloric ceca, with little to no involvement of the posterior intestine. This distribution pattern suggests regional susceptibility to the causative agent(s) along the intestinal tract. The standard cross-sectional sectioning of the intestine similarly demonstrated a variation in the severity of lesions, which suggests an inconsistent lesion distribution. This was consistent with the intra-fish evaluation of ten fish with ASE by Nervino (2022) on multiple cross sections taken along the length of the intestine. Nervino’s analysis was based on percent loss of epithelium, and five fish with moderate loss showed intra-fish variation ranging between 50-100% across individual cross sections. When included with the consistent distribution of lesions along the intestinal tract (as determined using Swiss rolls), these findings are likely influenced by the lesions being more prominent at the orad end, as well as some segmental distribution within the mid-intestine.

IHC analysis revealed distinct inflammatory patterns and tissue responses within ASE lesions. TGF-β showed positive labelling in individual to clustered cells within the mucosa and submucosa, suggesting 1) active cell growth, differentiation, and tissue repair, 2) attempts to dampen excessive inflammation, 3) a chronic versus acute process, and/or 4) the development of fibroplasia or early fibrosis. A chronic process with fibroplasia or early fibrosis is further supported by the development of granulation tissue in some ASE lesions. Previous publications in fish pathology have described granulation tissue as a component of granuloma complexes, particularly in bacterial infections (Miyazaki G Kaige, 1685; Mohi et al., 2010). Marty (1660) described “immature granulation tissue” as characterized by abundant vasculature, active fibroblasts, and collagen deposition. Our findings align with this definition; however, it is critical to distinguish granulation tissue from granulomatous inflammation and formation of granulomas, which are typically focal to multifocal and in response to a central infectious or foreign body nidus and poorly vascularized (Bruno et al., 2013; Pagán G Ramakrishnan, 2018). Chronic ASE lesions lack the organized epithelioid macrophage aggregates and encapsulation typical of granulomas. Instead, chronic ASE represents a diffuse, ulcerative enteritis with secondary granulation tissue formation as a wound-healing response. This distinction is important for accurate pathological classification and understanding of disease pathogenesis. Caspases are key driving enzymes in apoptosis, with caspase genes demonstrated in fishes (Gu et al., 2020; Zeng et al., 2021). Specifically, caspase-3 is involved in arresting the cell cycle and inactivating DNA repair, inactivating the inhibitors of apoptosis, and dismantling the cellular cytoskeleton (Nicholson, 1666). Caspase-3 is an intrinsic and extrinsic executioner caspase, so it is related to a very large infectious and non-infectious list of activators. This includes enteric protozoan parasite infections in mammals (Chin et al., 2002). In our study, caspase-3 labeling indicated ongoing apoptosis in damaged intestinal epithelial cells, while PCNA highlighted active cell proliferation in both regenerating epithelium and underlying tissues. We observed prominent caspase-3 activity in normal appearing gills, consistent with the physiologic epithelial turnover in gills (Sales et al., 2017), which may increase when injured (Topal et al., 2014). Together, these markers demonstrate that ASE lesions represent a dynamic balance between ongoing tissue damage, inflammation, and attempted repair.

Most samples for ASE descriptions and diagnosis in the present study as well as those of Nervino et al. (2024) and Nervino (2022) were from sexually mature broodstock fish that were humanely slaughtered and artificially spawned at hatcheries in the late summer. However, samples collected earlier in the run from the Sandy River Weir in 2023 and from earlier years in the Willamette River in the Nervino studies provided insights on the sequalae of intestinal lesions preceding the ASE lesions identified in spawned broodstock. These samples indicated that inflammation of the lamina propria precedes epithelial erosion, ulceration, and dysplastic change. Fish captured earlier in their river migration demonstrated an earlier stage of the disease (July, Sandy Wier), with a mean epithelial integrity of 87%, and ASE diagnosed in 6 out of 10 sampled fish. All but one fish had epithelial integrity scores of >80% epithelium, and amongst these fish the average inflammation score was 1.8, whereas Sandy Hatchery broodstock averaged 2.6. Severity of inflammation in the mucosa is associated with loss of the epithelium (Nervino et al., 2024). Nevertheless, consistent with the Sandy Weir fish collected in July, examination of the data from Willamette River fish collected mid-summer showed that 37% (n=25/67) of these fish had inflammation scores of 2 with > 90% epithelium intact. Additionally, the histologic presentation in recipient fish from the laboratory transmission study exposed to ASE by oral gavage showed a similar trend, with fish showing essentially intact, albeit dysplastic, epithelium with significant inflammation of the lamina propria in fish exposed to the ASE agent for only a few weeks (Polley et al., 2025). Thus, referencing the aforementioned pathogen-caused enteritis lesions, ASE is theorized to begin with leukocyte recruitment into the lamina propria, followed by erosion and ulceration of the epithelium, necrosis, and eventual granulation tissue as the body attempts to heal.

### 4.2 Parasite-ASE Associations

The relationship between parasitic infections and ASE remains a central question in understanding the disease etiology. This study examined the prevalence and distribution of two key parasites—*C. shasta* and *E. schreckii*—in relation to intestinal disease severity and geographic location of the sampled hosts. It is well recognized that enteric pathogens that damage the mucosa predispose the host to other infectious diseases. Examples in veterinary medicine include Parvovirus in dogs (Mylonakis et al., 2016), and relating closer to microparasites as seen here, there are numerous examples of coccidia exacerbating bacterial diseases in both poultry (Mesa-Pineda et al., 2021) and cattle (Sudhakara Reddy et al., 2013). Parasite infections in general often predispose the host to viral and bacterial diseases by general malaise or more specifically the phenomenon of diversion of immune response to the Th2 from the Th1, with the latter being more important for managing bacterial and viral diseases (Chen et al., 2023). This phenomenon also extends to teleost fishes (Okon et al., 2023). In salmonids, an example of another myxozoan that predisposes fish to bacterial diseases is *Tetracapsuloides bryosalmonae* (Hedrick et al., 1663). This includes exacerbated furunculosis in Coho Salmon (*Oncorhynchus kisutch*) that are naturally infected by the parasite (Higgins G Kent, 1666) and increased susceptibility to vibrio infections (Angelidis et al., 1687).

#### 4.2.1 Ceratonova shasta

*Ceratonova shasta* was the predominant parasite detected in this study, with an overall prevalence of 71.2% across all sampled broodstock populations. Notably, parasite prevalence was strongly associated with intestinal inflammation severity: among fish with severe inflammation (score 3), *C. shasta* prevalence reached 89.3%, compared to 0% in fish with no inflammation (score 0). This striking correlation raises the question of whether *C. shasta* is a driver of ASE or simply an opportunistic infection of damaged intestinal tissue. Nevertheless, *C. shasta* is not the primary cause of ASE. The parasite was absent in many fish with ASE in the multi-year retrospective study conducted by Nervino et al. (2024). A similar outcome was found in our present study, where in six endemic locations we did not detect *C. shasta* by histopathology in 12.4% (n=25/201) of the ASE positive fish. Moreover, the degree of epithelial loss amongst the *C. shasta* positive and negative fish was not statistically different, a critical indicator of ASE. *In vivo* laboratory transmission studies are very useful for elucidating roles of pathogens seen in field samples, which are numerous in adult Chinook Salmon (Polley et al., 2026). With ASE, it was shown that juvenile Chinook Salmon exposed to ASE tissues by oral gavage developed intestinal lesions consistent with the disease, but no *C. shasta* infections as it requires an intermediate host (Polley et al., 2025).

The geographic distribution of *C. shasta* also provides important insights. The parasite was completely absent from Puget Sound populations (White River and Minter Creek), which correspondingly displayed no ASE pathology. In contrast, Columbia Basin hatcheries exhibited very high prevalence of *C. shasta* across all locations (86%, n=185/215). This geographic pattern suggests that *C. shasta* exposure may exacerbate ASE development, potentially acting as a component, perhaps a promoter, of a multifactorial disease process with some fish.

#### 4.2.2 Enterocytozoon schreckii

The microsporidium *Enterocytozoon schreckii* so far has only been seen in fish with ASE (Couch et al., 2022; Nervino et al., 2024; Polley et al., 2025). It was detected in 20% of fish overall in the present study, with prevalence reaching 24.6% among fish with severe inflammation (i.e., ASE). Analysis of only positive locations showed a prevalence of 24.2% (n=54/223). However, this prevalence estimate is likely conservative due to a critical detection limitation; *E. schreckii* apparently infects only enterocytes, and its confirmatory identification requires the observation of mature spores within the cytoplasm of intact enterocytes, either mucosal or detached within the intestinal lumen. Hence, in fish with severe ASE, where epithelial loss can exceed 95%, the parasite may be present but undetectable due to loss of its host cell population.

Histological examination revealed that *E. schreckii* has a distinct anatomical distribution within the intestinal tract, predominantly infecting enterocytes in the pyloric ceca and mid-intestine of the intestinal tract. The parasite was specifically observed in healthy, attenuated but still-present epithelial cells, and detached luminal enterocytes. This anatomical preference overlaps with regions most severely affected by ASE lesions, raising questions about whether *E. schreckii* contributes to epithelial damage or simply exploits already-compromised cells. The prevalence of *E. schreckii* varied markedly among samples, with the highest detection at Willamette Hatchery (55.7%) and much lower rates at Round Butte (2.3%) and absent in fish from the Sandy Weir collected earlier in the summer. Interestingly, Willamette Hatchery broodstock retained relatively more epithelium (mean 57.6%) compared to Round Butte (3.2%) with only one or two enterocytes infected, which may explain the higher detection rate; more intact epithelium provides more opportunity for parasite visualization by histopathology. This detection bias must be considered when interpreting prevalence data and assessing the parasite’s role in ASE pathogenesis.

#### 4.2.3 Other Parasites

Larval cestodes were commonly observed in histological sections, with prevalence ranging from 5.4% to 80% across all hatcheries. Notably, cestode presence showed no association with ASE severity. Fish with heavy cestode burdens exhibited inflammation scores ranging from 0 to 3, but many fish with severe ASE had no detectable cestodes. This lack of association indicates that cestodes are incidental findings rather than contributors to ASE pathogenesis. Interestingly, the cestodes observed were marine larval stages. Our previous study (Polley et al., 2026), using wet mounts, identified this worm as a member of the genus *Pelichnibothrium*. The worm is identified by a scolex that has an apical sucker and four broadly attached bothridia, each with a small anterior, marginal accessory sucker, distinguishing it from other genera (Wardle G McLeod, 1652). The persistence of this marine parasite in the intestinal lumen of adult salmon after months in freshwater is a notable biological observation, demonstrating the longevity of these parasites in a larval state. Consistent with this, the Sandy Wier fish showed higher cestode prevalence than fish at spawning two months later. Likewise, the cestode was quite common in fish from both Puget Sound locations. Acanthocephalans in fish from Minter Creek were occasionally observed but were similarly not associated with ASE lesions, representing incidental parasites rather than disease contributors.

### 4.3 Geographic and Temporal Patterns

The geographic distribution of ASE reveals striking regional differences that provide critical insights into disease etiology and risk factors. Our expanded survey across the Columbia Basin and Puget Sound demonstrates that ASE is not a ubiquitous phenomenon in returning adult Chinook Salmon but rather is geographically restricted to specific watersheds.

#### 4.3.1 Puget Sound vs. Columbia Basin Broodstock

Based on examination of adult salmon from a limited number of rivers (Table 4), ASE appears to be a phenomenon linked to the Columbia River and tributaries. Complete lack of ASE or early lesions in all 44 broodstock fish from both Puget Sound hatcheries suggest that infectious agent(s) rather than natural, non-infectious progression of senescence in spring Chinook Salmon is the cause of the disease. Aside from the connection with Puget Sound, rather than the Columbia River, a distinction between these two hatcheries is that the distance from the ocean compared to ASE-affected locations is much shorter. Nevertheless, the summer water temperatures in White River range from 13-18 °C, similar to hatcheries in Oregon with ASE. Minter Creek holds fish in large tanks, but the same practice was used on some fish in Oregon that still exhibited ASE (Benda et al., 2015; Nervino et al., 2024). We have not detected *C. shasta* or *E. schreckii* at either Puget Sound hatchery, but it should be noted that based on histology many fish from ASE positive facilities are not infected with either parasite.

Whereas the Minter Creek fish showed no ASE, one fish exhibited profound proliferation of ECGs in the intestine. The cells appear as well-differentiated, mature ECGs, and hence a diagnosis of neoplasia was equivocal. The stratum granulosum of the gastro-intestinal tract of salmonids is replete with ECGs but may respond to antigenic stimulation in other layers. Kent et al. (1663) described unusual ECG proliferation in the intestines of many adult Coho Salmon (*O. kisutch*) from a hatchery in British Columbia, Canada. The lesions extended through all layers of the intestine and into the viscera. As we only examined intestines from Minter Creek, we could not assess the visceral distribution of the lesion in this fish in our present study.

#### 4.3.2 Within-Columbia Basin Variation

Even within the Columbia Basin, substantial variation in ASE severity was observed among sampling locations. Round Butte Hatchery had the most severe ASE across the four years of sampling, with the maximum mean present of only 7% across the groups. Carson Hatchery exhibited the next most severe pathology. Sandy Weir showed the mildest disease but was also a population sampled two months before spawning. Fish returning to different hatcheries traverse varying distances and encounter differing environmental parameters. Round Butte broodstock for example, often experience high water temperatures (as discussed below) and *C. shasta* exposure before transitioning to the cooler water hatchery environment. Hence, hatchery water sources, temperatures, and holding densities vary among facilities, all these environmental factors may influence disease progression, immune function, and parasite replication rates.

### 4.4 Implications for Prespawn Mortality and Temperature

Both ASE and elevated water temperature have been independently linked to PSM in adult Chinook Salmon (Benda et al., 2015; Carey et al., 2024; Nervino et al., 2024; Peterson et al., 2022), and these three factors are interactive. The severity of ASE pathology observed in this study—with some hatcheries showing mean epithelial retention as low as 3.2% (Round Butte) and inflammation scores of 3.0 (Carson and Round Butte)—indicates profound intestinal dysfunction that likely contributes to PSM. Although adult salmon in freshwater do not feed, such extensive mucosal damages face impaired osmoregulation, pathogen translocation through ulcerated surfaces, dysbiosis, potential increased risk of extraintestinal *C. shasta* infection, and systemic inflammation or shock that accelerates physiological decline.

A critical observation from multiple hatchery systems is that cooler water holding appears to reduce PSM without preventing ASE development. Benda et al. (2015) collected adult Chinook Salmon early in the freshwater run and held them at approximately 12°C in tanks fed by groundwater in two consecutive years. Nervino et al. (2024) compared ASE severity based on histopathology between these fish and their cohorts held in much warmer water at Willamette Hatchery. Whereas mortality was significantly reduced in cool water, the level of ASE based on percent epithelium present was identical in both populations within corresponding years. A similar pattern was seen in fish at Carson Hatchery, where fish are held at 8-10°C. According to hatchery staff, PSM is consistently low at this facility, while almost all the fish that we examined had ASE, and many had *C. shasta* and *E. schreckii* infections.

The Round Butte Hatchery also uses cold spring water (10°C) as its water supply, and here we observed the most severe ASE across the four sample years. However, PSM has been quite elevated in certain years at Round Butte despite always holding fish in cold water through the summer – e.g., 57% PSM in the first year of our study (2022), and 85% the year before. This location is unique, with an unusual high prevalence of renal ceratomyxosis, along with renomegaly and splenomegaly, indicating systemic dissemination beyond the intestinal tract (Polley et al., 2026).

This form of the infection was likely an important driver of PSM at this location. In contrast, Polley et al. (2026) provides an extensive review of pathogens in adult Pacific salmon, with discussion on links to PSM. A consensus of the authors, including fish health experts from California through Alaska, is that *C. shasta* is not an important cause of mortality in adult salmon, except when extraintestinal infections occur. Indeed, Benda et al. (2015) reported that successfully spawned Chinook Salmon broodstock from the Willamette River system were seven times more likely to have *C. shasta* than PSM fish.

The mechanism underlying the temperature effect of ASE-associated mortalities likely involves multiple factors. Cold water reduces metabolic rate (Brett G Glass, 1673), slowing pathological progression of PSM (Benda et al., 2015). Elevated temperatures often cause salmon to be more susceptible to infections; it not only enhances proliferation of pathogens but also may have immunosuppressive effects (Scharsack G Franke, 2022). Therefore, lower temperatures may also suppress efficaciousness of secondary infections in damaged tissues (Egan et al., 2025; Holt et al., 1686; Sanders et al., 1678). These observations indicate that while cold water does not prevent ASE development or parasite infection, it substantially slows clinical disease progression and extends the window for successful spawning. For hatchery management, these findings suggest that access to cold water holding facilities represents a practical but palliative intervention to reduce PSM and extend lifespan in broodstock populations with high ASE prevalence. The protective effect of cold water has concerning implications under climate change scenarios, particularly for wild fish that spawn in rivers. In addition, hatcheries currently benefiting from cold water sources may see those sources warm over time, eroding their advantage in broodstock survival. Adaptive management strategies accounting for changing thermal regimes will be essential for maintaining viable salmon populations under future climate scenarios.

### 4.5 Conclusion and Future Directions

Here we provided additional information on the geographic distribution of ASE and associated enteric pathogens, with examinations from 9 sampling locations (Table 4), adding to the survey by Nervino et al. (2024) that focused on the Willamette River and tributaries. This study demonstrates that ASE of Chinook Salmon is a geographically restricted, progressive intestinal disease strongly associated with *C. shasta* and *E. schreckii* infections in Columbia Basin. The complete absence of ASE in Puget Sound populations, combined with temporal progression data from the Sandy River system, argues against ASE as a normal senescent process and supports an infectious etiology. Detailed histopathological characterization defined ASE as a severe chronic, ulcerative and necrotizing enteritis.

Our survey should still be considered somewhat of a preliminary geographic survey, as there are still many spring Chinook Salmon populations from California through Alaska, and extending east to Idaho, that have yet to be examined. In addition to more locations, the potential occurrence of ASE in fall and winter run Chinook Salmon and other species should be examined. Multi-year sampling, or even ocean-run sampling of respective Columbia River cohorts, would reveal whether ASE prevalence and severity fluctuate with environmental conditions (e.g., river temperature, flow patterns) or remain relatively stable within populations.

Our use of histological examination for parasite detection has inherent limitations. *E. schreckii* prevalence is likely underestimated in fish with severe enteric ulceration, as the parasite requires intact enterocytes for visualization. Hence, one of us (C.S.) is developing as a specific qPCR test for the parasite. Regarding etiology focused on parasites, we did systematically investigate viral or bacterial pathogens. Polley et al. (2025) identified a putative viral pathogen in recipient juvenile fish with ASE-like lesions, suggesting that viruses may play a role in disease etiology. Hence, studies are underway to elucidate the presence of specific viruses and bacteria that are linked to ASE. For example, Couch et al. (2023) showed differences in the microbiome of fish based on severity of ASE from a small sample size. Microbiome work is in progress in Dr. H. Arnold’s program at Oregon State University using a much larger sample size of ASE positive and negative fish in attempt to identify microbial signatures of ASE.

## Acknowledgements

We thank Ryan Couture, Luke Whitman and Christopher Boyd with the Oregon Department of Fish and Wildlife, for assistance with collecting fish in Oregon. We also thank Emma Miller (Oregon Department of Fish and Wildlife) for providing *R. salmoninarum* ELISA. We thank Claire Couch, Connor Long, and Christian Serrano for assisting in sample collection in Oregon. We thank Dr. Charlene Morotti, Laura Swaim, Vannesa Ortiz, Matthew Stinson, the Northwest Indian Fish Commission, the Muckleshoot Indian Tribe, Joe Kalisch, and Washington Department of Fish and Wildlife for assistance collecting samples at White River and Minter Creek hatcheries and Larry Zeigenfuss for assistance with samples from Carson Hatchery in Washington state. We thank Sandra Garcia Maceiras at the University of Santiago de Compostela for providing excellent immunohistochemistry work.

## Data Availability Statement

The data that support the findings of this study are openly available at https://doi.org/10.7267/gt54kx16f (ScholarsArchive@OSU, Oregon State University).

