## Supplemental Figure 1-3 (S1-3) for "Geographic distribution of Adult Salmon Enteritis (ASE), *Enterocytozoon schreckii*, and *Ceratonova shasta* in Chinook Salmon *Oncorhynchus tshawytscha* (Walbaum, 1792) in Oregon and Washington state and expanded histopathologic description of ASE"

**9 Supplemental Figures**

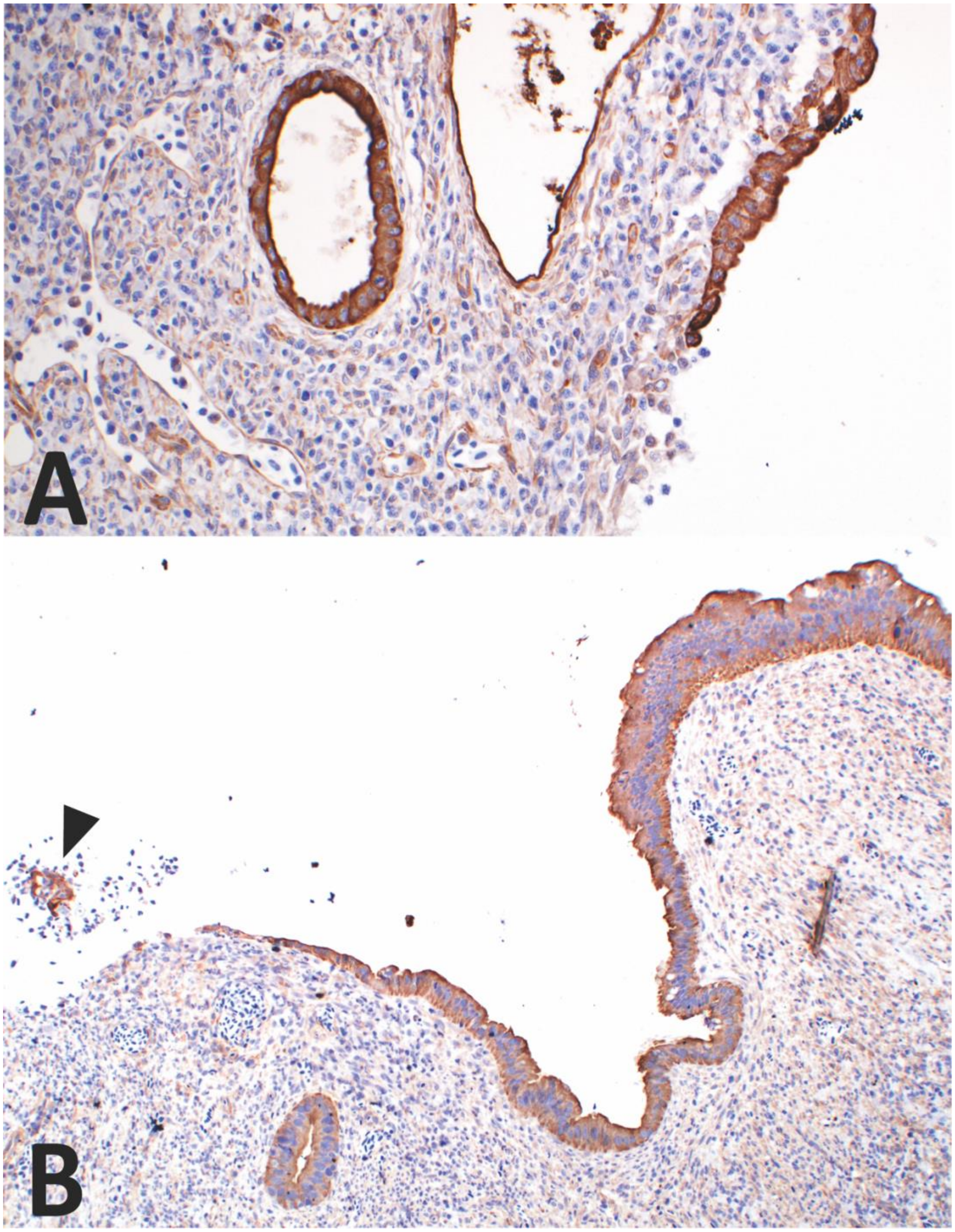


Figure S1. Characterization of the mucosal surface of adult salmon enteritis (ASE)-affected Chinook Salmon intestine using immunohistochemistry, where attenuated epithelial cells are distinguished from myofibroblasts by positive cytoplasmic uptake of cytokeratin WSS. A-B) Transition of intestinal mucosa from an erosive to an ulcerated lesion with attenuation of the enterocytes and increased cytokeratin uptake within detached luminal enterocytes (arrowhead).

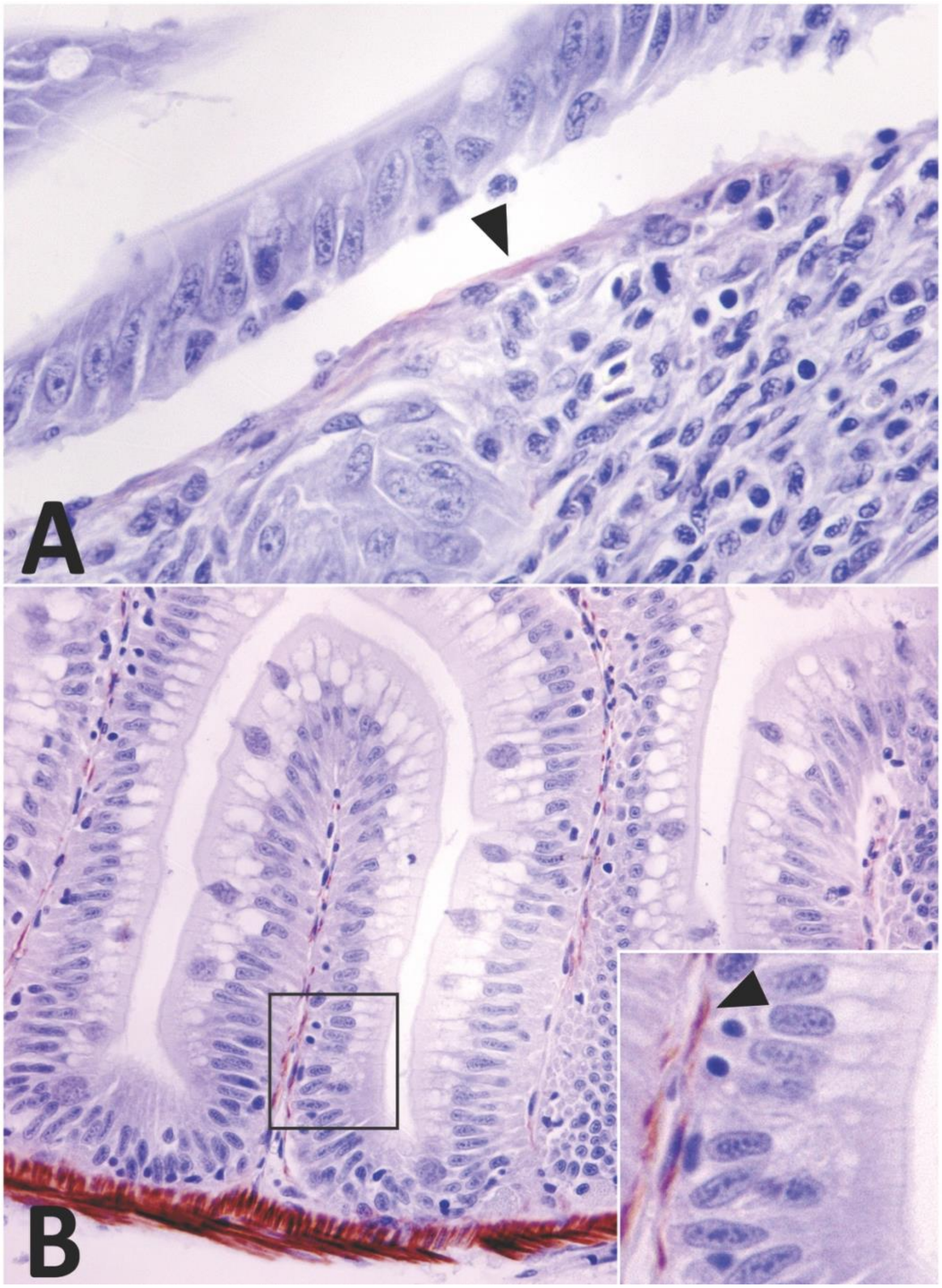


Figure S2. Immunohistochemistry for muscle actin-containing cells within the intestine of Chinook Salmon (A) and Zebrafish (*Danio rerio*) (B). A) Chinook salmon intestine affected by adult salmon enteritis. Subepithelial spindloid cells, consistent with myofibroblasts, have positive uptake for muscle actin (arrowhead). Space between epithelium and subepithelial cells is a processing artifact. A) Intestine from zebrafish, where subepithelial cells consistent with myofibroblasts stain positive (brown) for muscle actin. The smooth muscle of the muscularis shows strong uptake. Inset shows myofibroblast-like cells with positive uptake at higher power (arrowhead).

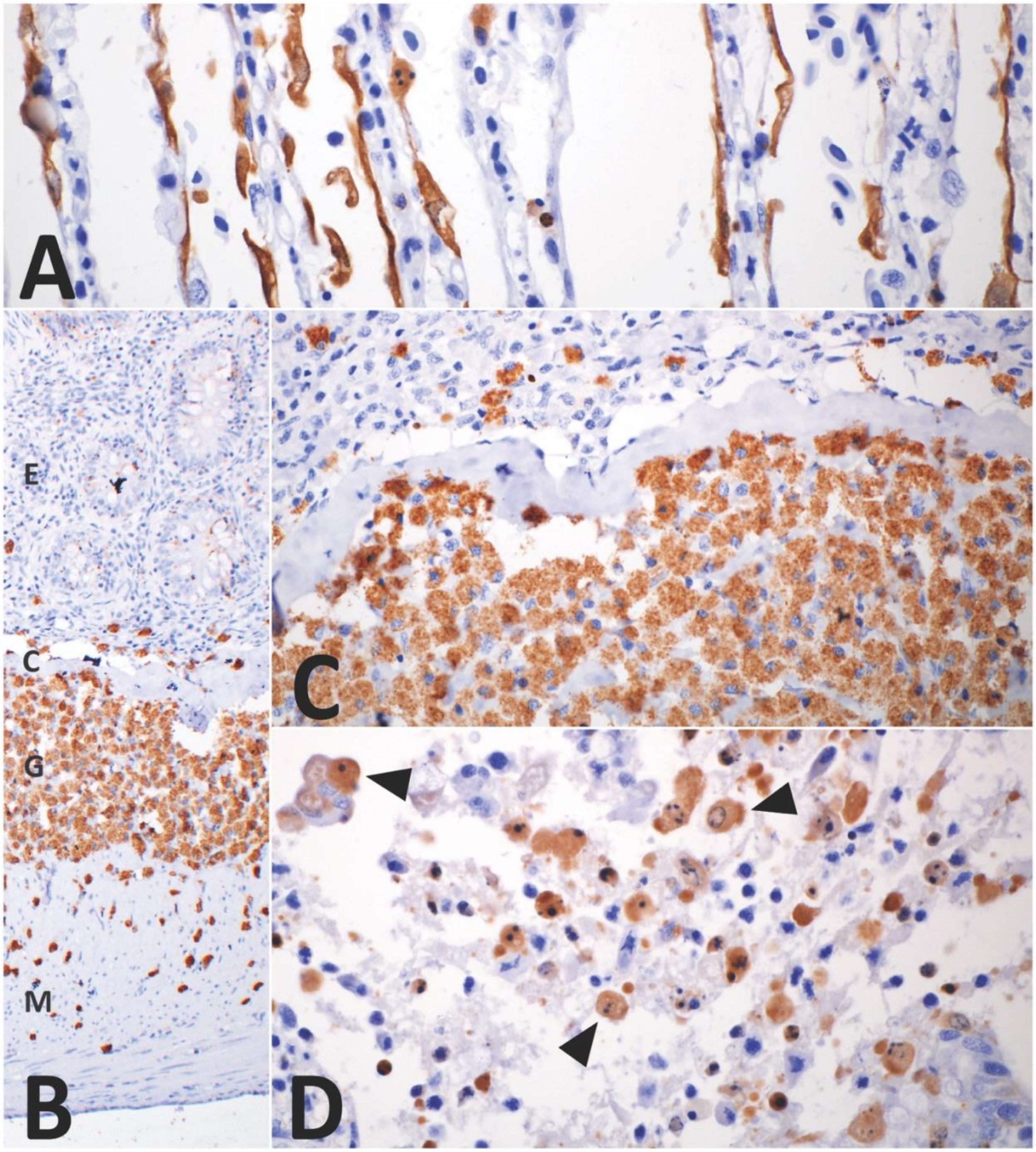


Figure S3. Immunohistochemistry for caspase-3, targeting apoptotic cells in Chinook Salmon gill (A) and intestine (B-D). A) Gill tissue with positive cytoplasmic uptake in gill epithelium. B) Intestinal section showing mucosa (E), stratum compactum (C), stratum granulosum (G), muscularis (M), and serosa with positive uptake in cells in the mucosa, stratum compactum, stratum granulosum, and muscularis. C) Cytoplasmic granules have strong uptake within granular cells within the stratum granulosum, with a few granular cells entering the overlying lamina propria. D) Luminal cellular debris, including detached enterocytes, from the mucosa (arrowheads), with strong positive cytoplasmic uptake.
